# DDHD2 provides a critical flux of saturated fatty acids to support neuronal energy demands

**DOI:** 10.1101/2023.12.31.573799

**Authors:** Saber H. Saber, Nyakuoy Yak, Xuan Ling Hilary Yong, Tobias Binder, Siyuan Lu, Reshinthine Purushothaman, Lidiia Koludarova, Safak Er, Irena Hlushchuk, Arnaud Gaudin, Tuula Nyman, Jeffrey R. Harmer, Gert Hoy Talbo, Mikko Airavaara, Brendan J. Battersby, Ashley J van Waardenberg, Victor Anggono, Giuseppe Balistreri, Merja Joensuu

## Abstract

Mitochondrial ATP production is fuelled by a fatty acid flux generated by phospholipase and triglyceride lipases in metabolically demanding tissues such as heart and liver, while the brain has long been believed to use almost solely glucose for energy. Phospholipase A1 enzyme DDHD2 is a major triglyceride lipase in the brain, and the loss of DDHD2 function results in a saturated free fatty acid (sFFA) imbalance and lipid droplet (LD) accumulation in the brain. The LD accumulation in neurons has been enigmatic as LDs are mainly considered to serve as a fuel storage. Here, we demonstrate that the loss of DDHD2 results in a mitochondrial respiratory dysfunction that leads to a significant decrease in ATP production and acetyl coenzyme A levels in neurons, even when the glycolytic breakdown of glycose occurs normally. Loss of DDHD2 also leads to a presynaptic defect as well as an imbalance in the global protein homeostasis in the neurons. These defects were rescued by external supplementation of the sFFA myristic acid coupled with its cofactor coenzyme A (Myr-CoA), indicating sFFA fuelling for neuronal β-oxidation. We have thus discovered that the sFFAs released by the activity of DDHD2 play a central role in providing energy to fuel synaptic function.

**One Sentence Summary:** Free fatty acids released by DDHD2 activity play a central role in maintaining neuronal energy levels and synaptic function.

## Introduction

DDHD2 is a mammalian intracellular phospholipase A1 that cleaves acyl ester bonds of phospholipid glycerol moieties from phospholipid membranes and from triglycerides in lipid droplets (LDs) (*1, 2*), generating saturated free fatty acids (sFFAs) and 2-acyl-lysophospholipids (*3–5*). Mutations in *DDHD2* gene, resulting in a loss of the protein domain responsible for membrane binding, as well as phospholipase and triglyceride hydrolase activities, cause hereditary spastic paraplegia 54 (HSP54), which is a childhood autosomal recessive neurodegenerative disorder characterized by neuromuscular and cognitive impairments (*6–15*). To date, the mechanisms by which *DDHD2* gene mutations cause HSP54 are not well understood, and at present, there is no cure or effective treatment for the disease.

Consistent with its role in lipid metabolism and in the central nervous system, the loss of DDHD2 function results in the accumulation of LDs in human HSP54 patient brains (*6*) and in mouse neurons (*1, 2*). As LDs are primarily considered a fuel storage, and the brain has long been believed to use almost solely glucose for energy, the observed LD accumulation in neurons following the loss of DDHD2 function has remained enigmatic. We recently discovered an activity-dependent release of saturated free fatty acids (sFFAs), in particular myristic acid (C14:0), palmitic acid (C16:0), and stearic acid (C18:0), *in vitro* in hippocampal neuron cultures, and *in vivo* in the brains of wildtype C57BL6/J mice in response to learning and memory formation (*5*). This activity-dependent release of sFFAs was blocked in *DDHD2* knockout mice (*1*) (*DDHD2^−/^)* following both *in vivo and in vitro* assays (*5*).

Given that sFFAs are an essential energy source and play important roles in regulating membrane trafficking and protein lipidation, here we investigated whether loss of DDHD2 function affected neuronal energy metabolism, protein homeostasis, membrane fluidity and synaptic membrane trafficking. We discovered significant reduction of acetyl coenzyme A levels in *DDHD2^−/−^* mice brain tissues, together with an impaired mitochondrial respiratory function and ATP production in cultured *DDHD2^−/−^* neurons. Importantly, the decreased ATP production in *DDHD2^−/−^* neurons was observed in the presence of glucose, and we observed no significant changes in the glycolytic capacity of *DDHD2^−/−^* neurons compared to control C57BL6/J neurons. The loss of DDHD2 function also led to an imbalance in global proteostasis and a substantial defect in synaptic vesicle recycling, marked by the accumulation of large bulk endosomes. The impaired mitochondrial respiratory function and ATP production were fully rescued within 48 h with an external supplementation of myristic acid coupled with its cofactor coenzyme A (Myr-CoA), suggesting Myr-CoA fuelling of β-oxidation in neurons. In support of functional b-oxidation in neurons, we demonstrate that cultured mouse cortical neurons express all the mitochondrial carnitine cycle and β-oxidation proteins required to shuttle long-chain FFAs into mitochondria at RNA and protein levels. External supplementation of Myr-CoA also led to a marked restoration of protein homeostasis, presynaptic function, and previously reported perturbations in secretory membrane trafficking (*16*), in cultured *DDHD2^−/−^* neurons, identifying Myr-CoA supplementation as a potential therapeutic intervention for HSP54. These unexpected findings suggest a role for DDHD2 in the synapse to provide a critical flux of sFFAs for mitochondrial β-oxidation in neurons, overturning the long held view that the brain uses glucose as its exclusive metabolic fuel (*17*).

## Results

### Impaired mitochondrial respiration and ATP production in *DDHD2*^−/−^ neurons are fully restored by myristic acid – CoA supplementation

Here, we examined how the loss of DDHD2 function affected mitochondrial function. First, using a fluorometric acetyl coenzyme A assay, we observed a significant decrease in the acetyl coenzyme A levels in *DDHD2^−/−^* brain tissues compared to wild-type control mice brain (Fig. 1A; 31.5% decrease compared to control). Acetyl coenzyme A is an essential cofactor necessary for the utilization of FFAs in the Krebs cycle, protein lipidation (N-myristoylation and S-palmitoylation), and as a fuel for energy production (i.e. β-oxidation) in mitochondria, among other functions.

**Fig. 1.**
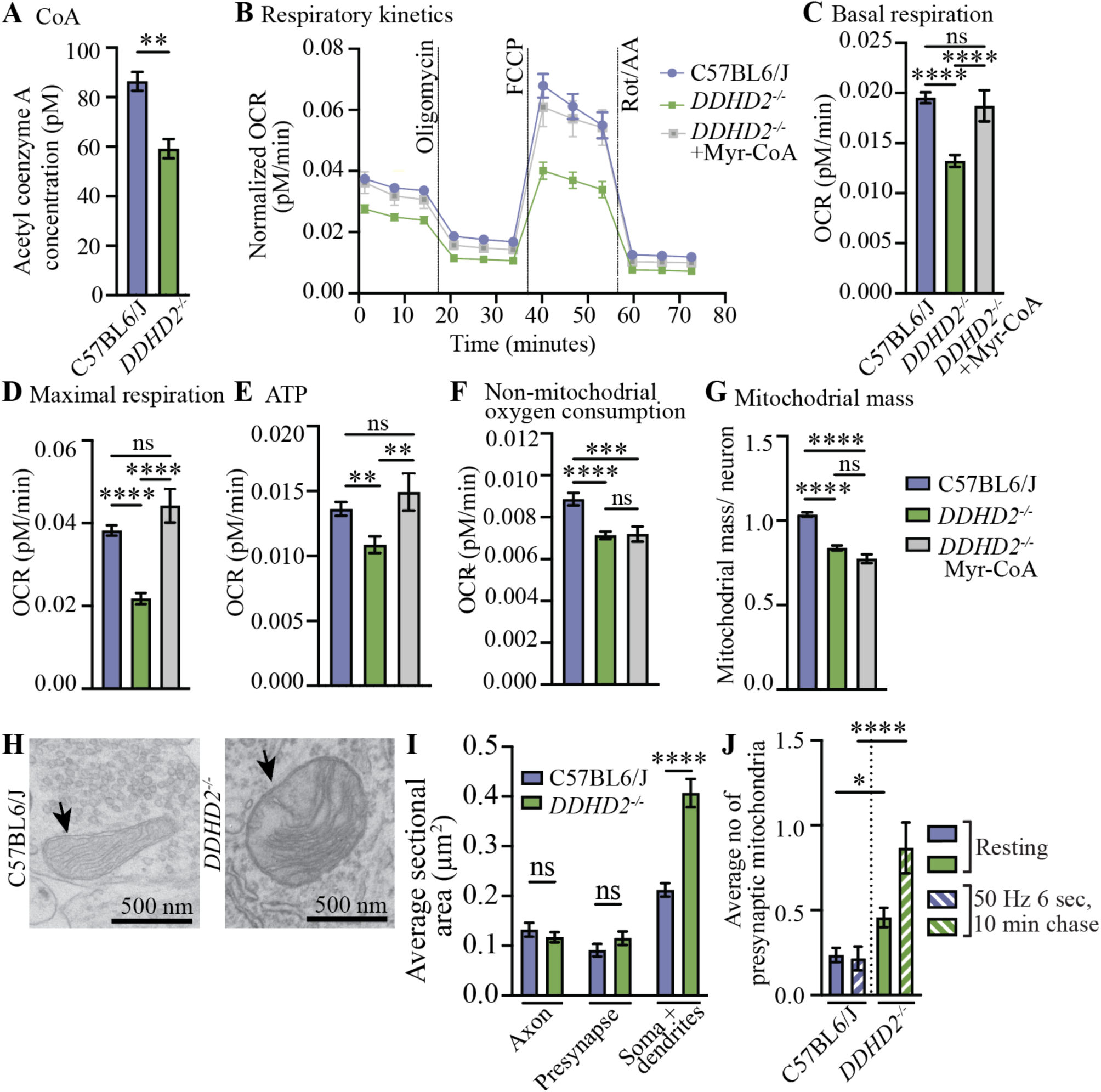
Myristoyl-coenzyme A restores mitochondrial respiratory function in *DDHD2^−/−^* neurons. (**A**) Fluorometric assay of acetyl coenzyme A (CoA) levels in C57BL6/J and *DDHD2^−/−^* brain lysates. (**B**) Seahorse XF oxygen consumption rate (OCR), (**C**) basal respiration, (**D**) maximal respiration, (**E**) ATP production, (**F**) non-mitochondrial oxygen consumption, and (**G**) mitochondrial mass measurements in cultured C57BL6/J, *DDHD2^−/−^* and *DDHD2^−/−^* neurons treated with 1 µM Myr-CoA for 48 h. (**H**) Representative electron microscopy (EM) images, and (**I**) quantification of average sectional area (µm^2^) of mitochondria (arrows) in cultured C57BL6/J and *DDHD2^−/−^* hippocampal neurons. (**J**) Quantification of number of mitochondria in cultured C57BL6/J and *DDHD2^−/−^* hippocampal neuron presynapses from EM images in resting (non-stimulated) condition and following 300 action potentials (50 Hz, 6 s) and 10 min chase at 37°C. Data are the mean ± sem (N=2 (A), N=3 (B-J) independent experiments). Student’s t test (A), Kruskal-Wallis (C-G) and 2way ANOVA with Tukey post hoc (I,J) multiple comparisons tests. Non-significant (ns), *p<0.05, **p<0.01, ***p<0.001, ****p<0.0001.

To test if the loss of function of DDHD2 lipase activity would affect neuronal mitochondrial function and energy levels, even in the presence of glucose, we assessed mitochondrial function using Seahorse Cell Mito Stress assay in control C57BL6/J and *DDHD2^−/−^* cultured neurons. The oxygen consumption rate (OCR) was measured as a direct readout for the mitochondrial respiration activity of the cells. We observed a significant OCR reduction in *DDHD2^−/−^* neurons compared to control neurons (Fig. 1B). Statistical analysis for each of the respiration stages showed in Figure 1E confirmed that compared to control neurons, *DDHD2^−/−^* neurons displayed significantly lower basal respiration (Fig. 1C), maximal respiration (Fig. 1D), ATP levels (Fig. 1E; 20.3% decrease), non-mitochondrial oxygen consumption (Fig. 1F), and mitochondrial mass (Fig. 1G). In striking contrast, the neuronal glycolytic capacity was unaffected by the loss of DDHD2 (Fig. S1A). Using Seahorse glycolytic rate assay (Fig. S1A), we observed similar basal respiration (Fig. S1B), glycolytic capacity (Fig. S1C), and non-glycolytic acidification levels (Fig. S1D) in control and *DDHD2^−/−^* hippocampal neurons, indicating that the decrease in ATP production was not due to a perturbed glucose metabolism.

We showed previously that loss of DDHD2 leads to a significant decrease in activity-dependent upregulation of sFFAs, in particular myristic acid (*5*). To be utilized as fuel in ß-oxidation, as a substrate for protein lipidation, or for the synthesis of complex lipids, sFFA must be first conjugated with coenzyme A (CoA). We found the levels of acetyl coenzyme A to be significantly lower in *DDHD2^−/−^* compared to control neurons (Fig. 1A). In addition, the coupling of sFFA with CoA requires ATP, which was also less abundant in *DDHD2^−/−^* neurons (Fig. 1E). We, therefore, tested if external supplementation of myristic acid preconjugated with CoA (Myr-CoA) would rescue the observed mitochondrial defects in *DDHD2^−/−^* neurons. Treatments with 1 µM Myr-CoA for 48 hours not only resulted in a complete rescue of the *DDHD2^−/−^* mitochondrial respiratory function (Fig. 1B-D) and ATP production (Fig. 1E), but it increased the ATP production by 9.5% compared to the levels in control neurons. In contrast, non-mitochondrial oxygen consumption was not affected by the supplementation (Fig. 1F), suggesting that Myr-CoA supplementation might specifically affect mitochondria function. Within the 48-hour rescue, we did not see an improvement in the mitochondrial mass, which might require prolonged treatment or supplementation with other sFFAs for a complete rescue (Fig. 1G). Notably, the conjugation of CoA with the myristic acid was essential because supplementation with 1 µM myristic acid without CoA failed to rescue mitochondrial respiratory function and ATP production (Fig. S2A-E). These results identify Myr-CoA as the minimal supplement to bypass the requirement for DDHD2 activity. Hence, we used Myr-CoA in our subsequent rescue experiments.

Neurons have long been considered to use almost solely glucose as their energy source. To test if neurons have the capacity for mitochondrial β-oxidation, we first investigated whether the β-oxidation protein machinery was expressed in cultured mouse cortical neurons. We found that all the factors required to shuttle FFAs into mitochondria and to oxidise them were robustly expressed at the RNA and protein levels as neurons mature and form synaptic connections (Fig. S3, and Table 1 and 2). Among the most upregulated proteins during synaptic maturation was Cpt1C (carnitine palmitoyltransferase 1C) (Fig. S3E), which converts fatty acyl-CoAs into fatty acylcarnitines that are transported into the mitochondrial intramembrane space through porins, the process of which is considered rate-limiting step for mitochondrial b-oxidation in times of energy requirement (*18–20*). This finding establishes that the enzymatic machinery to oxidize FFAs as substrates for mitochondrial aerobic ATP production is present in neurons.

### Loss of DDHD2 leads to an increase in the size of mitochondria in somatodendritic compartments and their accumulation in nerve terminals

To test if the observed defects in respiration corresponded to changes in mitochondrial ultrastructure in *DDHD2^−/−^* mice, we first performed electron microscopy (EM) analysis of mitochondria in cultured *DDHD2^−/−^* and controls hippocampal neurons (Fig. 1H). By quantifying the area of the mitochondria from EM thin sections, we discovered significantly enlarged mitochondria in the somatodendritic compartment of *DDHD2^−/−^* compared to controls, while no changes were observed in the average mitochondrial size in axons or presynapses (Fig. 1I). We observed a significant accumulation of mitochondria in resting (non-stimulated) *DDHD2^−/−^* presynapses compared to C57BL6/J controls. This difference increased following 50 Hz electrical stimuli (6 s, followed by 10 min incubation before fixation and processing for EM) (Fig. 1J). In support of these observations, our live cell imaging of mitochondria using MitoTracker dye in cultured C57BL6/J neurons treated with a pharmacological inhibition of N-myristoyltransferases (NMT) 1 and 2 (1 µM IMP-1088 for 48 h) (Fig. S4A; Movie 1 and 2) revealed that inhibition of protein N-myristoylation leads to a similar enlargement of mitochondrial size (Fig. S4B) and a reduction in the overall mitochondrial number (Fig. S4C) as observed in *DDHD2^−/−^* neurons. We also observed a significant reduction in axonal trafficking of mitochondria (Figs. S4A, S4D) following IMP-1088 treatment, and an accumulation of mitochondria in synapses. These results indicate that the sFFAs released by DDHD2 play a crucial role not only in the neuronal energy metabolism, but also in the structural integrity of mitochondria, and that the loss of DDHD2 function leads to a mitochondrial distribution defect in neurons.

### *DDHD2^−/−^* neurons have an altered proteome that can be significantly rebalanced by Myr-CoA supplementation

In addition to the mitochondrial dysfunction, we recently demonstrated that loss of DDHD2 leads to a perturbation in secretory pathway membrane trafficking (*5*). To characterize the broad-spectrum cellular changes arising from the loss of DDHD2 function further, we performed proteomic analysis by label-free mass spectrometry in cultured control and *DDHD2^−/−^* neurons. The effect of Myr-CoA supplementation on the entire proteome was also tested in this experiment. We identified 2,511 proteins that were quantified in at least 2 of the 3 replicates for each experimental group (Table 3). Replicate reproducibility of the analysis was assessed by principal component (Fig. S5A) and hierarchical clustering (Fig. S5B) analysis, confirming that the proteome identified in each replicate consistently clustered according to its respective experimental group (i.e., *DDHD2^−/−^*, C57BL6/J, and *DDHD2^−/−^* + Myr-CoA treatment).

Further analysis revealed a global proteomic shift in *DDHD2^−/−^* compared to control neurons (Fig. 2A, green dots). Of the 2,511 identified proteins, 965 proteins were significantly (adjusted p-value < 0.05) either up- or down-regulated (Fig. 2A, green dots). Supplementation with 1 µM Myr-CoA for 48 hours significantly reduced this difference, with more than 60% (605 proteins) of the 965 proteins that had altered expression levels in *DDHD2^−/−^* were no longer significantly different from control neurons (Fig. 2C).

**Fig. 2.**
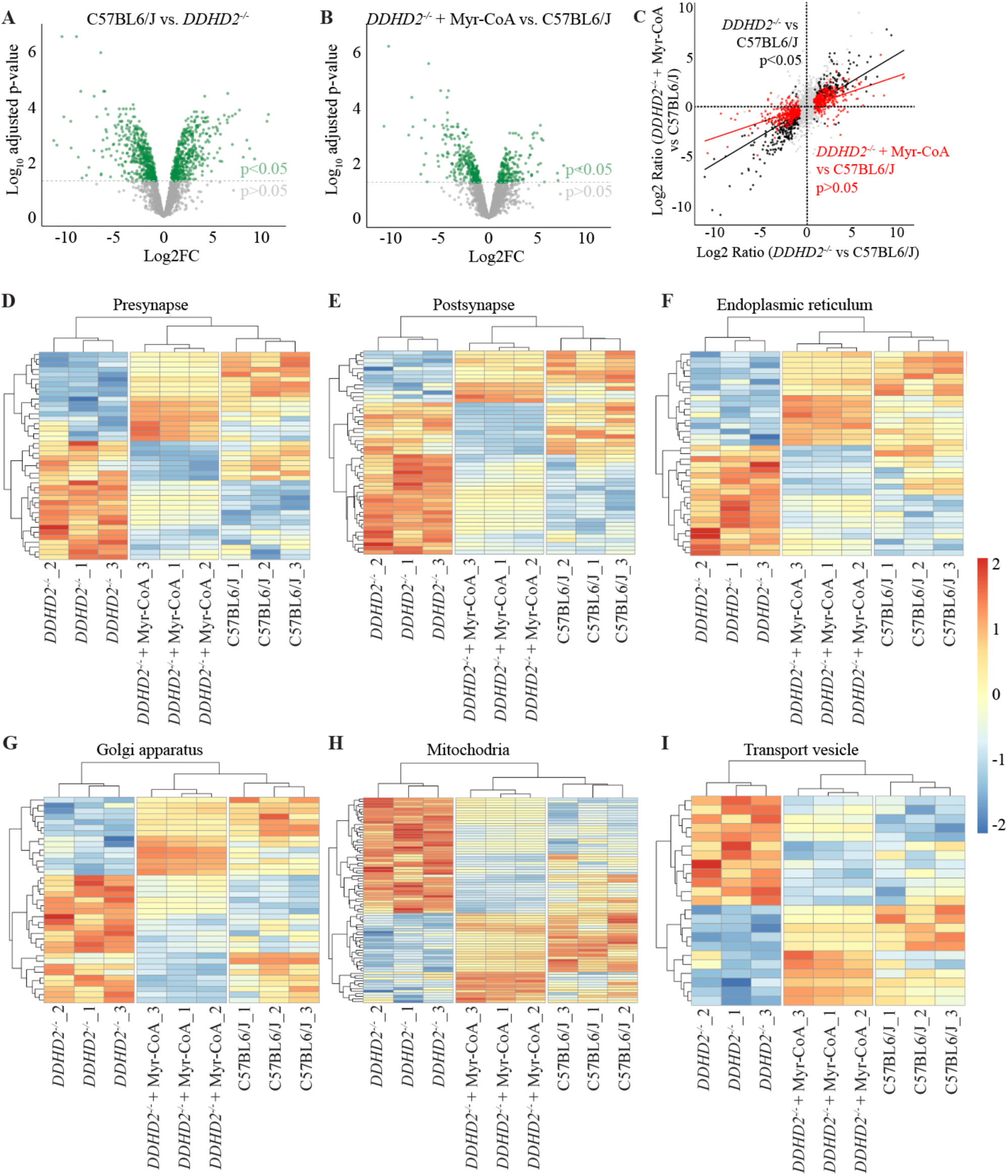
Altered *DDHD2^−/−^* hippocampal neuron proteome is partially rescued with Myr-CoA. Volcano plot of mass spectrometry proteomics of cultured (**A**) C57BL6/J vs. *DDHD2^−/−^* and (**B**) *DDHD2^−/−^* neurons treated with 1 µM Myr-CoA for 48 h vs. C57BL6/J. Altered protein abundance is represented as the log2 FC (or ratio) of the C57BL6/J vs. *DDHD2^−/−^* (**A**, X-axis) and *DDHD2^−/−^* +Myr-CoA vs. C57BL6/J (**B**, X-axis), and −log10 transformed adjusted p-value (Y-axes, p<0.05 threshold indicated as a dotted grey line, N=3 each condition). (**C**) Protein abundance of *DDHD2^−/−^* vs. C57BL6/J (X-axis) and *DDHD2^−/−^* +Myr-CoA vs. C57BL6/J (Y-axis). Proteins with unchanged levels (grey dots) and those significantly altered in abundance in *DDHD2^−/−^* vs C57BL6/J (965 proteins, black dots) are highlighted. Notably, 605 proteins (red dots) regain normal levels after Myr-CoA rescue in *DDHD2^−/−^* The black and red lines correspond to linear model fits to these sets of proteins, highlighting the *DDHD2^−/−^* proteome shift towards C57BL6/J. (**D-I**) Protein abundance heatmaps for indicated pathways in indicated conditions. Each row represents an individual protein. Cells are coloured by the z-score calculated for each row. Samples (columns) and proteins (rows) are clustered euclidean method with complete linkage.

Pathway analysis of proteins associated with pre- and postsynapse, endoplasmic reticulum, Golgi apparatus, as well as mitochondria and transport vesicles, demonstrated that, following Myr-CoA treatment, protein expression levels in each pathway were consistently more similar to the control than to the *DDHD2^−/−^* group (Fig. 2D-I). In *DDHD2^−/−^* neurons, large clusters of proteins either increased (warm colours) or decreased (cold colours) in abundance compared to control neurons. Myr-CoA treatment inverted these patterns to levels more comparable to control neurons. This striking inversion in protein abundance was observed in each analysed pathway (Fig. 2D-I). Together, these results demonstrate that a short Myr-CoA treatment was sufficient to bypass the requirement for DDHD2 activity and significantly rebalance the global *DDHD2^−/−^* proteome towards that of control neurons.

### Impaired protein trafficking in the secretory compartment of *DDHD2^−/−^* neurons is rescued by Myr-CoA supplementation

We recently demonstrated in cultured neurons that the loss of DDHD2 function leads to dilation and arrest of transport vesicles in the Endoplasmic Reticulum Golgi Intermediate Compartment (ERGIC) and Golgi complex (*5*). As a follow-up of our mass spectrometry results identifying changes in the abundance of secretory pathway and transport vesicle proteins in *DDHD2^−/−^* neurons, here, we used single-particle tracking photo-activated localization microscopy (sptPALM) (*21, 22*) to image mEos2-tagged (photoactivatable green-to-red fluorescent protein) ERGIC-53, which is a mannose-specific membrane lectin operating as a cargo receptor transporting glycoproteins between ER, ERGIC and Golgi, in live neurons (Fig. 3A). Our results revealed that the loss of DDHD2 leads to a significant decrease in the ERGIC-53-mEos2 trafficking in cultured neurons (Fig. 3B-E), supporting the observed defect in membrane trafficking (*5*). This mobility defect was fully rescued with an external supplementation of 1 µM Myr-CoA for 48 h (Fig. 3B-E).

**Fig 3.**
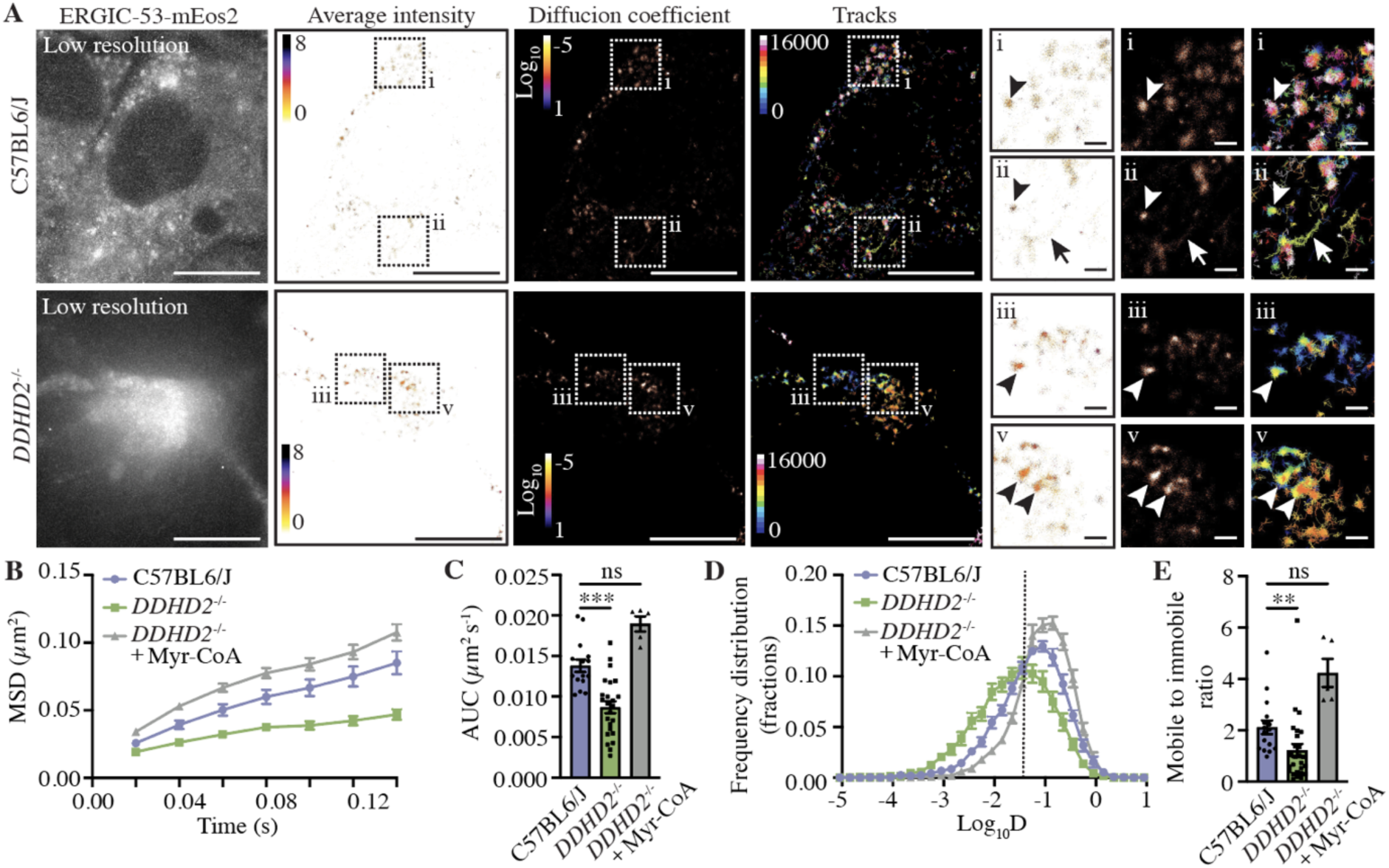
Super-resolution imaging of ERGIC-53. (**A**) sptPALM super-resolution imaging ERGIC-53-mEos2 in cultured C57BL6/J and *DDHD2^−/−^* neurons. Low resolution image of green fluorescence of the ERGIC-53-mEos2, along with super-resolved average intensity (bar: 8 to 0, high to low density), diffusion coefficient (bar: Log_10_ 1 to −5, high to low mobility), and single-molecule trajectory (bar: 0–10,000 frame acquisition) maps are show. Boxed areas (i-v) are shown magnified on right. Arrowheads point to confined molecules with low mobility, while arrows indicate mobile molecules. Quantification of single-molecule mobility of ERGIC-53-mEos2 in C57BL6/J and *DDHD2^−/−^* neurons (± 1 µM Myr-CoA for 48 h) shown as (**B**) mean square displacement (MSD), (**C**) area under the MSD curve, (**D**) frequency distribution of log_10_ diffusion coefficients ([D] = μm^2^ s^−1^), and (**E**) mobile to immobile ratio of diffusion coefficient frequency distributions (immobile Log_10_D ≤ −1.45 and mobile Log_10_D > −1.45). Data are the mean ± sem, Kruskal-Wallis multiple comparison test, N = 3 independent experiments, non-significant (ns), **p<0.01, ***p<0.001.

### Loss of DDHD2 perturbs the balance between synaptic vesicles recycling and activity-dependent bulk endocytosis

We next investigated how the loss of DDHD2 function affected the presynapses. The biogenesis and recycling of synaptic vesicles primarily rely on clathrin-mediated endocytosis (CME) at the presynaptic plasma membrane and endosomes (*23*). After endocytosis, internalized vesicles first fuse with and then are regenerated from presynaptic endosomes (*24–27*). To further explore the function of DDHD2 in synaptic vesicle recycling, we conducted quantitative EM analysis of hippocampal neurons derived from control and *DDHD2^−/−^* mice. Neurons were stimulated with high K^+^ buffer supplemented with the fluid phase endocytic marker horseradish peroxidase (HRP) for 5 min, after which the neurons were washed with low K^+^ buffer and chased for 10 or 30 min (Fig. 4A). Following the 10 or 30 min chase, we observed a significant decrease in the average size of HRP-positive endosomes in control neurons (Fig. 4A,B). At these timepoints, HRP positive synaptic vesicles were readily observed (Fig. 4A, white open arow heads). In contrast, the average size of endosomes in *DDHD2^−/−^* neurons increased significantly over the chase time (Fig. 4A,B). Quantification of the percentage of total HRP-stained endocytic structures revealed that the proportion of HRP-stained synaptic vesicles in control neurons was significantly higher than in *DDHD2^−/−^* neurons following both 10- and 30-min chase times, with the majority of the HRP signal remaining in large endosomal structures in *DDHD2^−/−^* neurons (Fig. 4C).

**Fig. 4.**
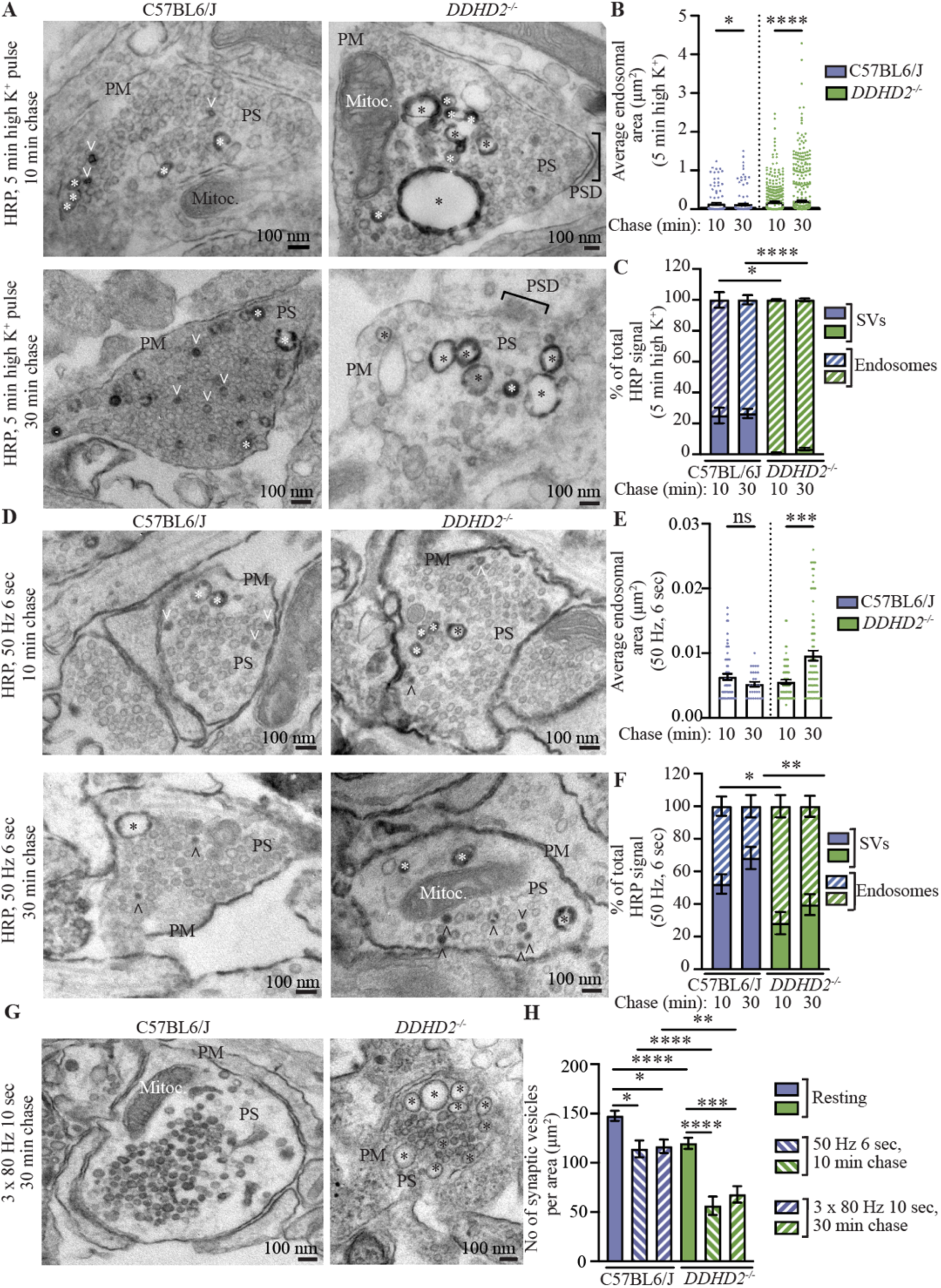
Loss of DDHD2 leads to a perturbed balance between synaptic vesicle recycling and activity dependent bulk endocytosis. (**A**) Representative electron microscopy (EM) images of cytochemically stained C57BL6/J and *DDHD2^−/−^* cultured hippocampal neuron presynapses (Ps) following a 5 min high K^+^ pulse with horse radish peroxidase (HRP) and either 10 min or 30 min chase at 37°C. Quantification of (**B**) average endosome area (µm^2^) and (**C**) percentage (%) of total HRP-stained synaptic vesicles (SVs) and endosomes in indicated conditions. (**D**) Representative EM images and quantification of (**E**) average endosome area (µm^2^) and (**F**) % of total HRP-stained SVs and endosomes in cytochemically stained cultured C57BL6/J and *DDHD2^−/−^* hippocampal neuron presynapses where HRP uptake was activated by 300 action potentials (50 Hz, 6 s), followed by either 10 min or 30 min chase at 37°C. (**G**) Representative EM images and (**H**) comparison of total number of SVs per presynaptic area (µm^2^) in resting (non-stimulated), 50 Hz, 6 s and 3×80 Hz, 10 s (800 action potentials) conditions in C57BL6/J and *DDHD2^−/−^* hippocampal neurons, and followed by either 10 min or 30 min chase at 37°C. Endosomes (asterisks), synaptic vesicles (SVs, open arrowheads), plasma membrane (PM), mitochondria (Mitoc.), and postsynaptic density (PSD) are indicated. Data are the mean ± sem (N=3 (A-C), N=1 (D-E)). Kruskal-Wallis multiple comparisons test. Ns, non-significant, *p<0.05, **p<0.01, ***p<0.001, ****p<0.0001.

To further support the results obtained with K^+^ stimulation, two different electrical field stimulation experiments were performed. Firstly, a train of 300 action potentials delivered at 50 Hz for 6 s in the presence of HRP was conducted (Fig. 4D). Confirming our earlier results, the average endosomal area decreased significantly from 10 min to 30 min chase in control neurons, consistent with efficient synaptic vesicle reformation. In contrast, *DDHD2^−/−^* neurons exhibited an increase in the average size of HRP-stained endosomes following the same chase time (Fig. 4E). Furthermore, the proportion of HRP-stained synaptic vesicles was significantly higher in control neurons compared with *DDHD2^−/−^* neurons following both 10- and 30-min chase (Fig. 4F). It is noteworthy that electrical field stimulation is a milder stimulation compared to the 5 min high K^+^ buffer treatment. Consistent with this difference, the high K^+^ stimulation induced larger compensatory bulk endosomes than the electric stimulation (Fig. 4B,E).

In the second, stronger, electrical field stimulation experiment, both control and *DDHD2^−/−^* neurons were subjected to three sets of 800 action potentials (3x 80 Hz 10 seconds, 800 action potentials, 30 min chase) in the presence of HRP, followed by EM analysis. Image quantification demonstrated a significant decrease in number of synaptic vesicles in *DDHD2^−/−^* neurons under resting, and in neurons challenged with both 50 Hz 6s or 3x 80 Hz 10 s stimulatory conditions (Fig. 4G,H). Together, these results indicate that loss of DDHD2 perturbs the balance between activity-dependent bulk endocytosis and synaptic vesicle recycling.

### Supplementation of Myr-CoA rescues *DDHD2^−/−^* synaptic vesicle recycling defects during high-frequency stimulation

The retrieval of endocytic membranes at synapses is linked with exocytosis, ensuring, on one hand fidelity of neurotransmission, and on the other the preservation of plasma membrane integrity and synaptic size (*25, 28*). To directly monitor synaptic vesicle recycling, we followed the trafficking of the vesicular glutamate transporter 1 (vGlut1) fused to the pH-sensitive green fluorescent protein (pHluorin) (vGlut1-pH (*29*)). The construct functions as a reporter of synaptic vesicle exo- and endocytosis and vesicular reacidification. Its fluorescence intensity is quenched within the acidic vesicle lumen and subsequently dequenched following synaptic vesicle fusion with the plasma membrane or neutral endosomes (*30*). To explore the role of DDHD2 in synaptic vesicle recycling, hippocampal control and *DDHD2^−/−^* neurons, transiently expressing vGlut1-pH, were subjected to high-frequency stimulation (50 Hz, 6 s) (Fig. 5A). Even before stimulation, the basal fluorescence intensity of the reporter vGlut1-pH was higher in *DDHD2^−/−^* neurons compared to control neurons (Fig. 5B, zero time point), indicating that at any given time the extent of vGlut1-pH localized in non-acidic compartments (i.e. plasma membrane and neutral early endosomes) was higher in *DDHD2^−/−^* neurons compared to controls. Following stimulation, the fusion of synaptic vesicles to the plasma membrane resulted in a rapid increase in the vGlut1-pH fluorescence signal that, in our experiments, reached a peak (indicated as 1) in approximately 10 seconds both in control and *DDHD2^−/−^* neurons (Fig. 5B). Thus, loss of DDHD2 did not impair secretion of synaptic vesicles. The compensatory retrieval of membranes via endocytosis, on the other hand, results in the re-internalization of vGlut1-pH molecules into acidic synaptic vesicles, which corresponds to a gradual decrease in the fluorescence signal of the reporter (Fig. 4B). Compared to control neurons, DDHD2 depleted cells displayed slower kinetics of vGlut1-pH delivery into acidic vesicles (Fig. 5B), with a corresponding significant increase in the time constant value τ (Fig. 5C).

**Fig. 5.**
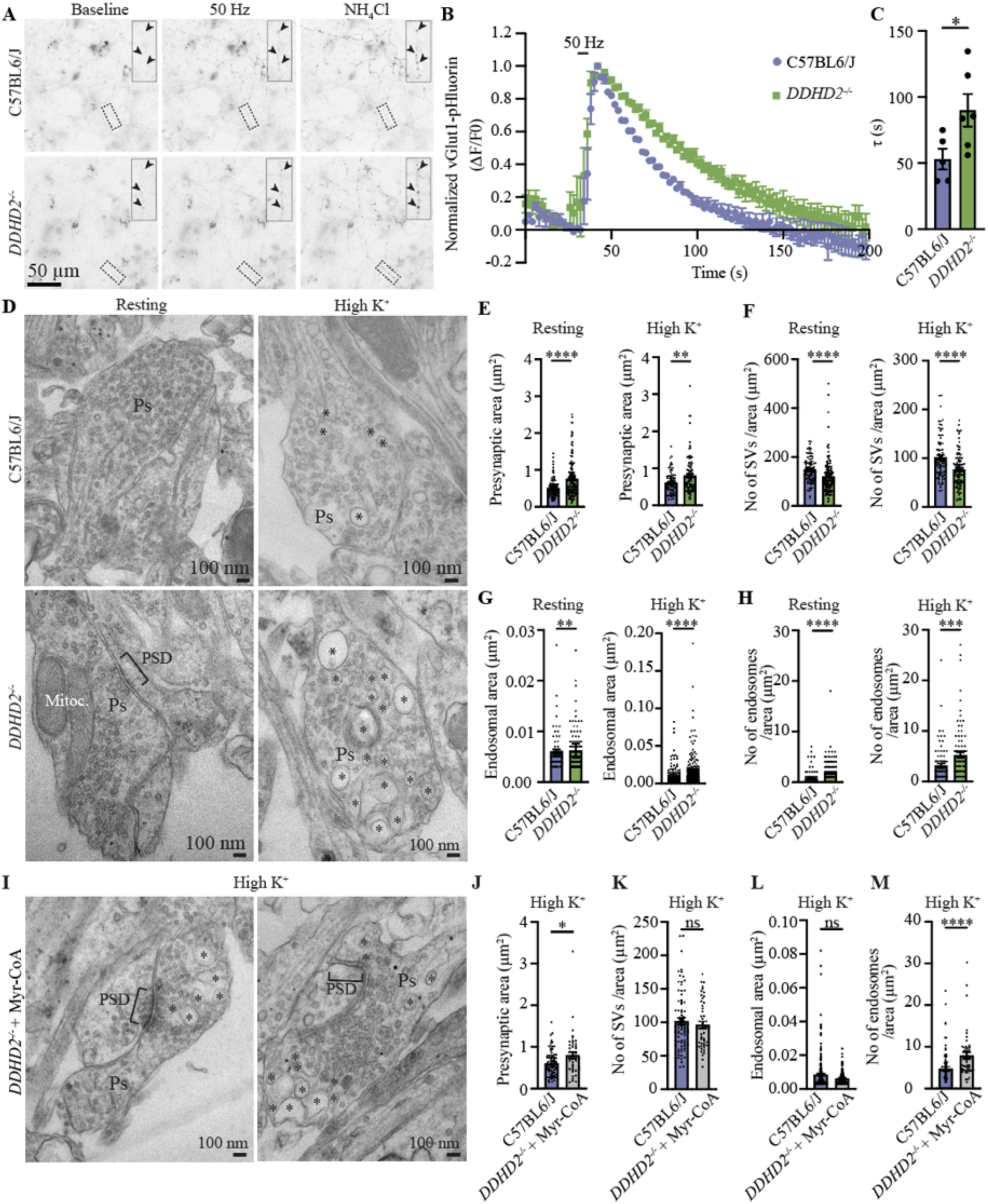
Loss of DDHD2 leads to perturbed synaptic vesicle recycling. (**A**) Representative widefield images of C57BL6/J and *DDHD2^−/−^* hippocampal neurons expressing vGlut1-pHluorin at rest (baseline), following 300 action potentials (50 Hz, 6 s), and NH_4_Cl dequenching. Boxed areas are magnified in the insets and arrowheads indicate presynapses. (**B**) Averaged, normalized vGlut1-pHluorin traces (ΔF/F_0_) and (**C**) endocytosis time-constant (τ) comparison in C57BL6/J and *DDHD2^−/−^* hippocampal neurons stimulated with a train of 300 action potentials (50 Hz, 6 s). (**D**) Representative electron microscopy (EM) images and quantification of (**E**) presynaptic area (µm^2^), (**F**) number of synaptic vesicles per presynaptic area (µm^2^), (**G**) endosome area (µm^2^), and (**H**) number of endosomes per presynaptic area (µm^2^) in cultured C57BL6/J and *DDHD2^−/−^* hippocampal neuron presynapses (Ps), in resting (non-stimulated) condition and following a 5 min high K^+^ stimulation. (**I**) Representative EM images and (**J-M**) quantification in cultured C57BL6/J, and *DDHD2^−/−^* hippocampal neurons rescued with 1 µM Myr-CoA for 16 h following a 5 min high K^+^ stimulation. Endosomes (asterisks), mitochondria (Mitoc.), and postsynaptic density (PSD) are indicated. Data are the mean ± sem (N=3), Student’s t test (C), and Mann–Whitney U test (E-H, J-M). Ns, non-significant, *p<0.05, **p<0.01, ***p<0.001, ****p<0.0001.

In support of our EM results demonstrating a deficiency in synaptic vesicle recycling in *DDHD2^−/−^* neurons, universal point accumulation in nanoscale topography (uPAINT) super-resolution imaging (*31*) of VAMP2-pHluorin (VAMP2-pH) bound Atto647N-nanobodies in live C57BL6/J and *DDHD2^−/−^* neurons in resting condition (low K^+^) and following high K^+^ stimulation (Fig. S7A) demonstrated that while we observed no significant changes in the nanoscale mobility of VAMP2-pH/Atto647N-nanobodies on the neuronal plasma membrane (Figs. S7B, S7C), the number of VAMP2 molecules on the plasma membrane was significantly reduced in *DDHD2^−/−^* neurons compared to control (Fig. S7D), supporting our earlier results demonstrating a perturbation in synaptic vesicle recycling.

To test if supplementation of sFFA could rescue these phenotypes, we included Myr-CoA in the neuronal growth medium and, following stimulation, used quantitative EM to monitor pre-synaptic area, number of synaptic vesicles, size and number of endosomes (Fig. 5I). This analysis revealed that a 16 h treatment with Myr-CoA was sufficient to decrease the overall presynaptic area in stimulated *DDHD2^−/−^* neurons, which, although still different, became closer to that of control neurons stimulated under the same condition (Fig. 5J). The number of synaptic vesicles per area was completely restored by the supplementation, becoming non statistically significantly different from that of stimulated control neurons (Fig. 5K). Endosomal area, which was significantly larger in stimulated *DDHD2^−/−^* neurons became indistinguishable from that of C57BL6/J control neurons (Fig. 5L), while the total number of smaller endosomes increased compared to non-supplemented *DDHD2^−/−^* neurons (Fig. 5M). Supporting these results, using immunofluorescence staining of tetrameric adaptor protein 2 (AP-2) complex, which connects the membrane and endocytic cargo receptors to the structural components of the clathrin coat (*32, 33*), we found that loss of DDHD2 was associated with a significant decrease in the abundance of endogenous AP-2, as well as mislocalization of AP-2 to the perinuclear space in the soma (Fig. S6). This phenotype was rescued with externally applied Myr-CoA, as demonstrated by a significant increase in the abundance of endogenous AP-2, and a restoration of the protein localization (Fig. S6). Taken together, these findings demonstrate that the activity of DDHD2 plays a crucial role in synaptic vesicle recycling and maintaining presynaptic integrity. These results indicated that the administration of Myr-CoA can facilitate restoring the balance between activity-dependent bulk endosomes and synaptic vesicle formation and support the notion that large bulk endosomes membranes are a source for synaptic vesicles reformation following high stimulation.

### DDHD2 activity regulates neuronal membrane fluidity

The balance between sFFAs and unsaturated FFAs (uFFAs) is tightly controlled in neurons to maintain normal cellular functions including intracellular trafficking and exocytosis^40^. This ratio is a critical factor in determining the molecular ‘packing’ of lipids which contributes to membranefluidity^41^. DDHD2 activity-dependent generation of sFFAs could potentially change the lipid composition of the biological membranes, which subsequently would change the membrane fluidity. Following our recent discovery of the sFFA imbalance in *DDHD2^−/−^* (*5*), we next investigated if this imbalance would alter membrane order. For this, we performed membrane fluidity measurements using Electron Paramagnetic Resonance (EPR) with the nitroxyl radical probes 5 doxylstearic (5-DSA) and 16 doxylstearic acid (16-DSA). Each has a nitroxide at the 5^th^ and 16^th^ carbon position of acyl-chain to measure the local fluidity near the protein/aqueous interface and near the hydrophobic protein cores, respectively. The EPR spectrum of a spin label embedded in the membrane indicates the dynamics (fluidity) of the lipid bilayer of the membrane. As a positive control, we treated neurosecretory PC12 cells with 1 µM cyclodextrin for 30 min, to extract cholesterol from cell membranes (*34*), increasing the membrane fluidity (Figs. S7E-S7G). To characterise the degree of fluidity we calculated a membrane order parameter *S* from the EPR spectra of both 5-DSA and 16-DSA as described before (*35*). Our results showed that cyclodextrin treatment significantly decreases the membrane order parameter *S* and thus increases the fluidity of the membrane compared to control cells. Next, we analysed the membrane fluidity in C57BL5/J and *DDHD2^−/−^* neurons (Figs. S7E, S7F). Analysis of the 5-DSA and 16-DSA spectra in C57BL5/J demonstrated that compared to resting conditions, following a 5-minute-high K^+^ stimulation, the plasma membrane of control neurons showed an increase in the local fluidity near the protein/aqueous interface (i.e. 5-DSA), as described by the decreased membrane order (S), while 16-DSA spectra indicated no significant change in the magnitude of the membrane fluidity near the hydrophobic membrane core (Fig. S7H). Differently from control neurons, quantification of 5-DSA and 16-DSA spectra in *DDHD2^−/−^* neurons showed no response to high K^+^ stimulation (Fig. S7I). Together, these results indicate that in addition to regulating neuronal energy levels and membrane trafficking, DDHD2 activity contributes to maintaining membrane biophysical integrity.

## Discussion

Our results point to a paradigm shift in the current understanding of neuronal energy production. Historically, neurons have been thought to use glucose as an energy source to fuel the high energy-demanding synaptic-activity. Our data indicate that sFFA are, in fact, an important source of energy to support neurotransmission, and their utilization as energy fuel, even in the presence of saturating glucose concentrations, accounts for approximately 20.3% of the neuronal ATP levels. Losing this energy supply due to loss of DDHD2 function, as it occurs in HSP54, leads to severe functional defects at synapses, secretory pathway compartments, mitochondria, as well as a disruption of the global proteostasis. Additionally, we found that in mouse cortical neurons all the factors required to shuttle FFAs into mitochondria and to oxidise them were expressed at the RNA and protein levels as neurons mature and form synaptic connections. Similarly, recent proteomic studies on mitochondria purified from cerebellar neurons and astrocytes confirmed the presence of the b-oxidation machinery (*36*) and demonstrated that, albeit at lower levels compared to astrocytes, neuronal mitochondria performed efficient b-oxidation (*36*). In conjunction to our study, compelling evidence support this unexpected role of sFFA as an additional energy source in neurons (*37*).

We found that *DDHD2^−/−^* neurons have disturbed mitochondrial size, distribution, and functions, and cells have decreased levels of acetyl coenzyme A and ATP, even in the presence of glucose. We also found that in the absence of DDHD2, the entire proteome is significantly different from that of control neurons. We have demonstrated here that this plethora of defects can be rescued by the exogenous supplementation of a sFFA, myristic acid, only when coupled to CoA but not if administered alone. In addition to activating sFFA for mitochondrial utilization and as a substrate for protein lipidation, the coupling with CoA rendered the molecule readily soluble in aqueous solution, which is a favourable aspect for potential prophylactic or therapeutic use in HSP54.

The quantitative studies presented here link the loss of DDHD2 function to specific defects in synaptic membrane trafficking, particularly synaptic vesicle recycling and the turnover of bulk endosomes. Our result point to a defect in CME. These defects were markedly restored with the Myr-CoA supplementation, indicating that additional sFFAs, such as CoA-conjugated palmitic acid, may also be required for a full rescue of synaptic function. The molecular mechanism behind the CME-ADBE imbalance in *DDHD2^−/−^* neurons remains to be investigated in future studies.

Based on our results, we envision that the supplementation of a combination of sFFAs with CoA could provide a successful avenue for therapeutic intervention for pathologies linked to DDHD2 loss of function such as HSP54. In addition, given the unforeseen role of sFFA as a supplementary energy source in neurons, such metabolic supplementation approaches could aid brain function in conditions where glucose utilization becomes less efficient, as it is the case during aging.

## Acknowledgements

We thank the facilities and staff at The University of Queensland (UQ) Centre for Microscopy and Microanalysis (CMM) and the Queensland Brain Institute’s Advanced Microscopy Facility, and the support from the Queensland Node of Metabolomics and Proteomics Australia for their assistance with metabolomics/proteomics data acquisition. We also thank the UQ’s Protein Expression facility (PEF) and staff for their assistance with cloning the ERGIC-53-mEos2. We extend our gratitude to Adel Kechlar, Corey Butler and Jean-Baptiste Sibarita for kindly providing the PALMTracer Software.

## Author Contributions

M.J. conceived and designed the study, and supervised S.H.S., N.Y., S.L., T.B., and R.P. S.H.S performed the mitochondrial respiratory function assays, mass spectrometry and electric field stimulation with help from G.H.T, J.H., X.L.H.Y. and V.A. M.J. performed the EM and S.L. assisted with EM quantification. A.v.W performed the hippocampal proteomics data analyses. L.K., S.E., I.H., T.N., M.A. and B.J.B performed the cortical neuron experiments. G.B. participated in experimental planning and data analysis. A.G. helped with data analysis. N.Y., R.P. and T.B. performed primary tissue culturing and cell culturing, and image analysis. S.H.S, M.J., G.B. wrote the manuscript. All authors participated in proof-reading the manuscript and approved it for publication.

## Data and materials availability

The unique constructs created in this study will be deposited to Addgene upon acceptance of the manuscript to publication. All original data created in this study will be uploaded to The University of Queensland Research Data Management repository. Any additional information required to reanalyse the data reported in this paper is available from the lead author upon request.

## Material and methods

### Ethics and mouse strains

All experimental procedures using animals were conducted under the guidelines of the Australian Code of Practice for the Care and Use of Animals for Scientific purposes and were approved by the University of Queensland Animal Ethics Committee (2022/AE000770). DDHD2 knockout (*DDHD2^−/−^*) mice (*1*) in C57BL6/J background were sourced from the Scripps Research Institute in the United States. The mice were maintained in a 12-h light/dark cycle (80% intensity) at 18−24°C (30–70% room humidity) and housed in a PC2 facility with ad libitum access to food and water. Wildtype NMRI were obtained from Charles River and kept at the University of Helsinki animal facility under license KEK21-012.

### Primary embryonic hippocampal neuron cultures

Primary hippocampal neurons were obtained from C57BL6/J and *DDHD2^−/−^* at embryonic day (E) 16. Isolated hippocampi were prepared and cultured as previously described (*38*). Neurons were seeded at 100,000 neurons per 78.54 mm^2^ onto poly-L-lysine (Sigma-Aldrich, #P2636) coated dishes: 1 mg/mL PLL for glass bottom dishes (CellVis, #D29-20-1.5-N), and 0.1 mg/mL for plastic dishes (for EM). All experiments were performed at days in vitro (DIV) 21-22, unless otherwise stated, to ensure mature and functional synaptic connections.

### Primary embryonic cortical neuron cultures

Pregnant wildtype NMRI (Charles River) mice were sacrificed via CO_2_ and cervical dislocation. Hysterectomy was done to collect E15-17 mice embryos. Cortices from embryos were dissected as previously described (*39*), then pooled to isolate primary neurons. Briefly, dissected cortices were collected in a standard 15-mL canonical tube, washed thrice with Hank’s Balanced Salt Solution (HBSS) without Ca^2+^, Mg^2+^ (Gibco, #14175-053), incubated on ice for 10 min, semi-digested with trypsin (MP, #103139) for 15 min at 37°C. To isolate individual cells, trituration was done once with HBSS with 10% fetal bovine serum (FBS) (Gibco, #10500056) and DNAse I (Roche, #11284932001), then twice with only HBSS with 10% FBS. Cells were centrifuged at 0.8 rpm (1 min), resuspended, and centrifuged again at 0.4 rpm (30 s) in culture media. Supernatant was collected in a new canonical tube, centrifuged at 0.8 rpm (2 min) for the final washing step, then resuspended with media. Isolated cells were maintained in Neurobasal media (Gibco, #21103049), containing 1×B27 Supplement (Gibco, #17504-044), 500 µM L-glutamine (Gibco, #25030–032), and 100 µg/mL Primocin (Invitrogen, ant-pm). Cells were plated on pre-coated (0.01% poly-L-lysine; Bio-Techne Cultrex, #3438-100-01), 6-well (1-2 x 10^6^ cells/well) or 10 cm (14.6 x 10^6^ cells/plate) plates, then kept at 5% CO_2_, 37°C. Half medium changes were done every 3-4 days until the sample collection.

### Generation of pmEOS2-C1-ERGIC53 plasmid

The pmEOS2-C1-ERGIC53 plasmid was constructed by amplifying a PCR fragment of ERGIC53 of the pMXs-IP spGFP-ERGIC53 (addgene Plasmid #38270) using BamHI and NheI restriction sites from the pMXs-IP spGFP-ERGIC53 plasmid with the following primers: 5’-GACATCggatccGCTACCGGACTCAGATCT-3’ and 5’-GACGCTgctagcTATGATCTAGAGTCGCGG-3’. This PCR fragment was digested with BamHI and NheI restriction endonucleases then ligated to the pmEos2-C1 plasmid backbone digested with XhoI and BamHI restriction endonucleases. The pmEOS2-C1-ERGIC53 plasmid was constructed using using the In-Fusion® Snap Assembly Cloning Kit (Takara) and transformed into OmniMAX competent cells. Overnight cultures of the positive clones were grown in LB Kanamycin (30 μg/mL) media and plasmid DNA was extracted using the QIAprep Spin Miniprep Kit (Qiagen). Purified plasmid DNA was sequenced using ABI BigDye Terminator v3.1 Sequencing at the Australian with the following primers; forward primer: 5’ – CTGTACAAGTCCGGACTCAGATCTATGGACGAGCTGTACAAGGG – 3’, reverse primer: 5’ – TTATCTAGATCCGGTGGATCC-TCA-AAAGAATTTTTTGGCAGCTGC – 3’ at the Genome Research Facility (AGRF). Data analysis was performed using the software SnapGene® 5.3.

### RNA isolation, Illumina sequencing and analysis

Total RNA was isolated from NMRI mouse cortical neuron cultures at DIV1 and DIV20 using the Reliaprep RNA Miniprep system (Promega) followed by deoxyribonuclease I (NEB) treatment and subsequently purified using Agencourt RNAClean XP magnetic beads (Beckman Coulter). Illumina sequencing and analysis was performed at Azenta Life Sciences. Briefly, each library preparation was strand specific with polyA selection with ERCC spike-in followed by deep sequencing on an Illumina NovaSeq 2X150 bp with 22 to 51 million total reads per sample. Sequencing reads were trimmed with Trimmomatic v.0.36 and then mapped to the Mus musculus GRCm38 reference genome using STAR aligner v.2.5.2b. Differential expression between DIV20 and DIV1 was calculated using DESeq2 and the Wald test.

### Electron microscopy analysis and quantification

Electron microscopy (EM) analysis was performed on cultured C57BL6/J, *DDHD2^−/−^* (± 1 µM Myr-C0A for 48 h) hippocampal neurons. For structural analysis of mitochondria and presynapses, neurons were stimulated for 5 min in high K^+^ buffer (56 mM KCl (Ajax Finechem Pty Limited, #1206119), 0.5 mM ascorbic acid (Sigma-Aldrich, #A5960), 0.1% bovine serum albumin (BSA; Sigma-Aldrich, #A8022), 15 mM HEPES (Sigma-Aldrich, #H3375), 5.6 mM D-glucose (AMRESCO, #0188), 95 mM NaCl (Sigma-Aldrich, #S9888), 0.5 mM MgCl_2_ (Chem-Supply, #MA029), and 2.2 mM CaCl_2_ (Sigma-Aldrich, #C5080) at pH 7.4, 290–310 mOsm), and then fixed with 2 % glutaraldehyde (Electron Microscopy Sciences, #16210) in 0.1 M sodium cacodylate (Sigma-Aldrich, #C0250) buffer pH 7.4 for 20 min at room temperature, washed three time for 3 min with 0.1 M sodium cacodylate buffer, and processed for EM using standard protocols. Control samples were incubated in low K^+^ buffer (0.5 mM MgCl_2_, 2.2 mM CaCl_2_, 5.6 mM KCl, 145 mM NaCl, 5.6 mM D-glucose, 0.5 mM ascorbic acid, 0.1% (wt/vol) BSA and 15 mM HEPES, pH 7.4, 290–310 mOsm) (i.e. resting condition) for 5 min and then fixed as described above. For peroxidase cytochemistry, cultured hippocampal neurons were stimulated with high K^+^ buffer supplemented with either 10 mg/mL (high K^+^ stimulation assays) or 1 mg/mL (electric field stimulation assays) horse radish peroxidase (HRP; Thermo Fisher Scientific, #31490) for 5 min at 37°C in a 5% CO_2_ atmosphere, washed with low K^+^ buffer (with total volume of 8–10 mL) and chased for either 10 or 30 min at 37°C in a 5% CO_2_ atmosphere. For electric field stimulation, cultured neurons were challenged with 300 (50 Hz 6 s) or 800 (3 x 80 Hz, 10 s) action potentials, and chased for 10 or 30 min at 37°C in a 5% CO_2_ atmosphere. Neurons were then fixed with 2% glutaraldehyde and 2% paraformaldehyde (Electron Microscopy Sciences, #15710) in 0.1 M sodium cacodylate buffer, pH 7.4, for 20 min at room temperature, washed three times for 3 min with 0.1 M sodium cacodylate buffer, and processed for 3,3′-diaminobenzidine tetrahydrochloride (DAB; Sigma-Aldrich, #D5905) and hydrogen peroxidase (H_2_O_2_) cytochemistry using standard protocols. All samples were contrasted with 1 % osmium tetroxide and 2 % uranyl acetate before dehydration and embedded in LX-112 resin using BioWave tissue processing system (Pelco) as previously described (*40*). Thin sections (80-90 nm) were cut using an ultramicrotome (Leica Biosystems, UC6FCS) and were imaged with a transmission electron microscope (JEOL USA, Inc. model 1101) equipped with cooled charge-coupled device camera (Olympus; Morada CCD Camera). Images were acquired randomly. The quantification of the number synaptic vesicles, the number and size (sectional area, µm^2^) endosomes, presynaptic area, mitochondrial size (sectional area, µm^2^) in presynapses, axons and somatodendritic compartment, and the number of presynaptic mitochondria were quantified manually using ImageJ/Fiji (*41*) (https://imagej.nih.gov/ij/) and Adobe Photoshop (Adobe, 22.4.3 release) Count Tool. Vesicles with sectional area ≤0.002 µm^2^ were classified as synaptic vesicles, and those >0.002 µm^2^ as endosomes.

### Metabolic fluxes in primary neurons

To assesses metabolic fluxes, we used Seahorse XFe96 respirometry assay as previously described (*42*). The test utilizes the integrated injection ports on XF sensor cartridges to introduce respiratory modulators into the cell well during the assay, unveiling essential mitochondrial function parameters. We monitored the glycolytic flux and mitochondrial respiration in real time by measuring the extracellular acidification rate (ECAR) and oxygen consumption rate (OCR). Briefly, Primary hippocampal from C57BL6/J and *DDHD2^−/−^* were plated in precoated poly-L-lysine coated (1 mg/ mL) XFe96 multi-well cell plate at density of 25,000 cell/well. The key parameters of glycolytic function including basal glycolysis, glycolytic capacity and non-glycolytic acidification were assessed using a Seahorse XF glycolysis assay (Agilent Technologies, Berlin, Germany) according to the manufacturer’s instructions. Prior to the assay, XF sensor cartridges were hydrated and calibrated with Seahorse Bioscience calibrant overnight at 37°C incubator without CO_2_. At DIV 19-21, neurons were washed with prewarmed XF assay media DMEM without phenol red, sodium bicarbonate and sodium pyruvate for glycolysis or with DMEM with 10 mM glucose and 2 mM glutamine and without phenol red, sodium bicarbonate and sodium pyruvate for Mito stress. Neurons were then preincubated in assay medium (the same media) for 40 min at 37 °C without CO_2_. All the glycolysis modulators 20 μl of 60 mM glucose, 20 μl of 7 μM oligomycin, and 20 μl of 1.2 M 2-deoxy-D-glucose (2DG) were sequentially injected in port A, B, and C, while for mitochondrial respiration modulators, 20 μl of 6 μM oligomycin, 20 μl of 14 μM carbonyl cyanide p-trifluoromethoxyphenylhydrazone (FCCP), and 20 μl of 8 μM rotenone/antimycin A (Rot/AA) were consecutively injected in port A, B, and C.

### Fluorometric measurements of Acetyl CoA

20 mg of C57BL6/J and *DDHD2^−/−^* brain tissue samples were frozen rapidly in liquid N_2_ and pulverized. Acetyl Coenzyme A (CoA) levels in the brain tissues were analysed using Acetyl-Coenzyme Assay Kit (Sigma-Aldrich, #MAK039-1KT) following manufacturer’s instructions. Quantification of the CoA levels were performed against standard curve according to manufacturer’s instructions.

### Live imaging and of Mitochondria with Mitotracker

E16 C57BL6/J neurons cells were cultured on glass-bottom dishes as previously described(*43*). On DIV18, neurons were treated with DMSO (vehicle control) or 1 µM IMP-1088 for 48 h at 37 °C in 5% CO_2_. On DIV 20, neurons were incubated in 200 nM Mitochondria dye Mitotracker-488 (Millipore, cat. no. SCT136) diluted in low K^+^ buffer for 15 min at 37 °C in 5% CO_2_, then imaged using a Zeiss Plan Apochromat 63x/1.4 NA oil-immersion objective on a confocal/two-photon laser-scanning microscope (LSM 710; Carl Zeiss Pty Ltd, Australia) built around an Axio Observer Z1 body (Carl Zeiss) and equipped with two internal gallium arsenide phosphide (GaAsP) photomultiplier tubes (PMTs) and three normal PMTs for epi-(descanned) detection and two external GaAsP PMTs for non-descanned detection in two-photon imaging, and controlled by Zeiss Zen Black software. The number and size of mitochondria was quantified using ImageJ/Fiji Analyse Particle Tool as described previously (*44*). The quantification of metachronal mobility was manually analysed in 200×200 px axonal areas, counting the number of transported mitochondria during 50 s acquisition (2 frames s^−1^ imaging rate). N = 3 independent neuronal cultures in each condition.

### Proteomics analysis using high resolution Orbitrap mass spectrometry

The LC/MS/MS identification and quantification of the digested peptides from primary hippocampal neurons from C57BL6/J, *DDHD2^−/−^* and *DDHD2^−/−^* treated with 1 µM Myristoyl coenzyme A lithium salt (Myr-CoA, Sigma-Aldrich, # M4414) were performed using a S-Trap^TM^ (Protifi, NY) Micro Spin Column Digestion protocol. Briefly, samples were solubilised by adding 50 µL of S-Trap lysis buffer (10% sodium dodecyl sulphate (SDS) in 100 mM Tris, pH 8.0) to 50 µL of sample, before reducing by adding 20 mM of dithiothreitol (DTT) and heating at 70°C for 60 min. Cysteine residues were alkylated to prevent disulphide bond reformation using 40 mM iodoacetamide for 30 min at room temperature in the dark. 2.5 µL of 12% phosphoric acid was added, followed by 165 µL of S-Trap binding buffer (90% methanol in 100 mM Tris) to the acidified lysate. The sample mix was then centrifuged through the S-Trap column at 4,000 x g for 1 min followed by three washes with 150 µL S-Trap binding buffer, with 4,000 x g centrifugation between each wash. Peptide digestion was initiated by adding 25 µL of 50 mM ammonium bicarbonate buffer (pH 8) containing 2 µg trypsin (Sequencing Grade Modified Trypsin, Promega, # V5117) directly on top of the column and incubating overnight at 37 °C. Peptides were eluted by three successive aliquots of 40 µL of 5%, 50%, and 75% acetonitrile in 0.1% formic acid, respectively. Eluted peptides were dried down using a vacuum concentrator (Concentrator Plus, Eppendorf). Samples were redissolved in 20 µL of 5% ACN (aq) and 2 µl were injected to a trap column (Thermo 22 mm x 300 µm, 5 µm, C18) at a flow rate of 10µL/min. Following 3 min wash the trap column was switched in-line with a resolving column (Water nanoEase 100 mm x 150 µm, 1.8 µm, 100 Å). The samples were eluted by a gradient was held constant at 8% for 4 min, then was increased linearly to 24% at 47 min, to 40% at 53 min and to 95% at 57 min. The gradient held constant for 1 min, before returning to start condition at 8% over 1 min. Mass spectrometry using LC-MS/MS was performed using Ultimate UHPLC system coupled to an Exploris 480 mass spectrometer with a FAIMS Pro interface (Thermo Fisher Scientific) The FAIMS compensation voltages were −45 and −65 V. The electrospray voltage was 2.2 kV in positive-ion mode, and the ion transfer tube temperature was 295°C. Full MS scans were acquired in the Orbitrap mass analyser over the range of m/z 340-1110 with a mass resolution of 120,000. The automatic gain control (AGC) target value was set at ’Standard’, and the maximum accumulation time was ‘Auto’ for the MS. The MS/MS ions were measured in 6 windows from mass 350-470, in 18 windows from mass 465-645 and 5 windows from mass 640-1100 with an overlap of 1 m/z and quadrupole isolation mode. Analysis of data were performed using Spectronaut against a reference proteome with a Q-value cut-off of 0.05.

### Additional processing and statistical analysis of label-free mass-spectrometry

The standard, non-normalised output of Spectronaut (BGS Factory Report text file) was imported into R version 4.05 for further processing using a modified method described previously (*45*). For proteins that mapped to multiple annotations, the ‘best annotation’ method previously described was used (*45*). Each sample group was permitted to have up to one missing value (that is, an intensity not reported one or three individual replicates). Protein intensities of the parent group protein ‘PG.ProteinGroups’ were then log2 transformed and globally normalised using the quantile normalisation method (*46*). Missing values were assumed to be missing at random and imputed using the knn nearest neighbour averaging method (*47*). Following, unwanted sources of technical variation were removed by surrogate variable analysis (*48*). Sample clustering was confirmed using unsupervised principal component analysis and hierarchical clustering. Principle Component Analysis of the first two principal components using singular value decomposition, as implemented in pcaMethods version 1.82.0 and hierarchical clustering using ‘ward’ method for clustering distance ‘euclidean’. For hierarchical clustering, probabilities of clustering were determining using 100 bootstrap replications, as implemented in pvcluster version 2.2-0, and probabilities of the branch positions shown in the plot, together with the probabilities of clusters (red boxes). For differential protein abundance, generalised linear modelling with Bayes shrinkage as implemented in limma version 3.46.0 was performed and proteins were considered differentially abundant at a corrected p-value of 0.05 unless specified otherwise (adjusted for false discovery rate using the Benjamini and Hochberg method) (*49*).

### Gene Set Enrichment Analyses

Heatmaps for proteins belonging to a gene set (gene ontology term) were independently generated by extracting all Gene Symbols from ‘org.Mm.eg.db’ version 3.1.2 that matched a specific GO term of interest. This was then filtered by proteins that were differentially abundant between DDHD2KO myristic treated (nonstim) versus DDHD2KO (nonstim) (or comparison of interest). The z-score of the protein intensities across all samples (row-wise) was then calculated across samples and plot as a heatmap using the pheatmap version 1.0.12. Clustering distance was ‘euclidean’ and clustering method ‘complete’ linkage. Targeted pathways selected included: ‘Mitochondria (GO:0005739)’, ‘Presynapse (GO:0098793)’, ‘the endoplasmic reticulum (GO:0005783)’, ‘endoplasmic reticulum–Golgi intermediate compartment (GO:0005793)’, ‘ER to Golgi transport vesicle (GO:0030134)’, ‘Golgi apparatus (GO:0005794)’, ‘secretory granule (GO:0030141)’, ‘transport vesicle (GO:0030133)’, and ‘ribosome (GO:0005840)’.

### Quantitative mass-spectrometry analysis of mouse cortical neurons

The proteins were precipitated on amine beads as previously described (*50*). The precipitated proteins on beads were dissolved in 50 mM ammonium bicarbonate, reduced, alkylated, and digested with trypsin (1:50 enzyme: protein ratio; Promega, United States) at 37°C overnight. The resulting peptide mixture was purified by STAGE-TIP method using a C18 resin disk (3M Empore) before the samples were analysed by a nanoLC-MS/MS using nanoElute coupled to timsTOF PRO2 (Bruker) with 60 min separation gradient and 25cm Aurora C18 column. MS raw files were submitted to MaxQuant software version 2.4.7.0 for protein identification and label-free quantification. Carbamidomethyl (C) was set as a fixed modification and acetyl (protein N-term), carbamyl (N-term) and oxidation (M) were set as variable modifications. First search peptide tolerance of 20 ppm and main search error 10 ppm were used. Trypsin without proline restriction enzyme option was used, with two allowed miscleavages. The minimal unique + razor peptides number was set to 1, and the allowed FDR was 0.01 (1%) for peptide and protein identification. Label-free quantitation was employed with default settings. UniProt database with ‘mouse’ entries (2020) was used for the database searches. Additional data filtering and statistical analysis was done using Perseus ver 1.6.1.5. using normalised intensities (LFQ).

### Fluorescence imaging of vGlut1-pHluorin trafficking

To monitor synaptic vesicle recycling, we utilized the pH-sensitive green fluorescent protein (pHluorin) fused with vesicular glutamate transporter 1, vGlut1-pH (*29*). The construct functions as a reporter of synaptic vesicle exo- and endocytosis and vesicular reacidification. Its fluorescence intensity is quenched within the acidic vesicle lumen and subsequently dequenched following synaptic vesicle fusion (*30*). Live hippocampal C57BL6/J and *DDHD2^−/−^* neurons transiently expressing vGlut1-pHluorin were mounted in an imaging chamber with field stimulation electrodes (RC-21BRFS; Warner Instruments, Hamden, CT) and continuously perfused with imaging buffer (119 mM NaCl, 2.5 mM KCl, 2 mM CaCl_2_, 2 mM MgCl_2_, 25 mM HEPES, 30 mM D-glucose, pH 7.4) supplemented with 10 μM NBQX (Abcam, #ab120046) and 50 μM DL-APV (Abcam, #ab120271). All experiments were performed at 35°C. The imaging solution was kept constant at 35°C using an inline solution heater (SH-27B; Warner Instruments). Before stimulation, the basal fluorescence intensity of the reporter vGlut1-pHluorin was recorded for in each C57BL6/J and *DDHD2^−/−^* acquisition. Neurons expressing vGlut1-pHluorin were challenged with a train of 300 action potentials delivered at 50 Hz, 6 s (100 mA and 1 ms pulse width) and imaged at 0.5 Hz (2 × 2 binning) through a 40X (1.4 NA) oil objective using an inverted Zeiss Axio Observer Z1 epifluorescence microscope equipped with an Andor Luca R EMCCD camera. At the end of each imaging acquisition, neurons were perfused with an alkaline imaging buffer (50 mM NH_4_Cl substituted for 50 mM NaCl) to reveal total pHluorin expression. Equal-sized regions of interest were placed over nerve terminals to measure the pHluorin fluorescence elicited by stimulation over time using the Time Series Analyzer plugin in FIJI software (NIH). Activity-dependent changes in fluorescence (ΔF/F0) were normalized to the respective peak heights from the train of stimuli (to calculate the rate of endocytosis) or to the total amount of fluorescence present after alkaline treatment (to calculate the exocytosis amplitude). The rapid increase in the vGlut1-pHluorin fluorescence signal following stimulation represents fusion of synaptic vesicles to the plasma membrane, while the compensatory retrieval of membranes via endocytosis results in the re-internalization of vGlut1-pHluorin molecules into acidic synaptic vesicles, which corresponds to a gradual decrease in the fluorescence signal of the reporter. The rate of fluorescence decline is described by the endocytosis time constant (τ value) (i.e. the time required for the fluorescence signal to reach zero if the rate of the decline was linear) which was calculated by fitting the decay phase for each trace to a single exponential function (*29*).

### Immunofluorescence

Rescue experiments with 1 µM Myr-CoA for 16 h were performed on cultured *DDHD2^−/−^* neurons by adding the supplement directly into the culture media. Neurons were then fixed in 4% paraformaldehyde in PBS for 30 min at room temperature, washed in PBS, permeabilized with 0.1% Triton x100 (Sigma-Aldrich-Merck, #X100-100ML) for 4 min and blocked with 1% BSA in PBS for 30 min. Neurons were then immunostained against endogenous AP-2 using Anti-AP-2 complex subunit alpha-1 (Abcam, #ab218624) diluted in blocking solution (1% BSA in PBS) over night at 4 °C, and detected using secondary antibody donkey anti-goat Alexa Fluor® 647 (Thermo Fisher Scientific, #A21447) was diluted in blocking solution (1% BSA in PBS) for 45 min at room temperature. Neurons were stained with DAPI (Sigma-Aldrich, #D9542) and Phalloidin Atto647N (Sigma-Aldrich, #65906) according to manufacturer’s protocols. Finally, neurons were washed and mounted in ProLong Gold Antifade Mountant (Thermo Fisher Scientific, #P36934). Confocal images were acquired with a spinning-disk confocal system consisting of an Axio Observer Z1 equipped with a CSU-W1 spinning-disk head, ORCA-Flash4.0 v2 sCMOS camera, 63x 1.3 NA C-Apo objective. Image acquisition was performed using SlideBook 6.0. Sections of each slide were acquired using a Z-stack with a step of 100 nm. The exposure time for each channel was kept constant across all imaging sessions. Images were deconvolved on Huygens deconvolution software 21.10 (Scientific Volume Imaging). For the quantification of AP-2 levels in C57BL6/J and *DDHD2^−/−^* neurons, fluorescence images acquired by confocal microscopy using identical microscope settings where first Z-projected using the Image-J software (Fiji version). The quantification of AP2 fluorescence was then performed using the Cell profiler 4 software (www.cellprofiler.org). First the nuclei (Primary Object) of the neurons were selected using the “Identify Primary Object” tool with the Otzu algorithm inbuilt in the software. Apoptotic nuclei, which were significantly smaller than healthy nuclei, were excluded from the analysis by automatic size exclusion. The perimeter of the cell cytoplasm was then automatically detected using the AP-2 fluorescence channel by using the “Identify secondary object” tool of the software with the Otzu algorithm and a very sensitive threshold of segmentation so that the whole cell contour was accurately outlines even in cells with very low AP-2 levels. From this ‘secondary object’, the nuclear area was excluded using the “Identify Tertiary Object” tool of the software. The resulting area of the ‘tertiary object’ was used to quantify the levels of AP-2 per cell. Visual inspection was performed for each image to make sure the segmentation of the cell cytoplasm was accurate. Obtained values where then background subtracted and normalized to the intensity of fluorescence of the respective nuclei stained with the nuclear dye Hoechst.

### Electron paramagnetic resonance measurements (EPR)

Analysis of biophysical EPR parameters fluidity measurement were carried out as previously reported (*51*). To assess membrane fluidity, the following nitroxyl radicals SLFA probes were used: Spin-labelled stearic acids2-(3-carboxypropyl)-4,4-dimethyl-2-tridecyl-3-oxazolidinyloxy (5-doxyl-stearic acid, 5-DSA, cat. no.253618) and 2-(14-carboxytetradecyl)-2-ethyl-4,4-di methyl-3-oxazolidinyloxy (16-doxyl-stearicacid, 16-DSA, cat. no. 253596) from Sigma-Aldrich. 5-DSA (spin probe containing the nitroxide group attached on C5 that is located on the opposite terminal relative to the charged carboxyl fatty acid terminus) was used to determine the local fluidity near the protein/aqueous interface, and 16-DSA (spin probe containing the nitroxide group attached on C16 that is located on the opposite terminal relative to the charged carboxyl fatty acid terminus) was used to assess the fluidity near the hydrophobic protein cores. C57BL6/J and *DDHD2^−/−^* neurons, and neurosecretory PC12 cells were labelled 0.02 M methanol-containing solutions of 5-DSA or 16-DSA and loaded into the capillary tubes to measure the electron paramagnetic resonance (EPR) spectra at 37°C using a Bruker E540 Benchtop Magnettech MiniScope MS5000 spectrometer (now Bruker Biospin, Berlin) equipped with bio temperature control and computerized data acquisition and analysis capabilities. Instrumental parameters for these measurements were: microwave frequency 9.47 86 GHz, microwave power 5.024 mW (16 dB attenuation of 200 mW source) 0 mW, modulation frequency 100 kHz, modulation amplitude 0.12 mT (16-DSA) or 0.3 mT (5-DSA). Resonances occur in the approximate, and magnetic field range 332346–342 356 mT. The number of scans varied depending upon the required signal-to-noise ratio; each scan was 60 seconds, and the measurement time was typically 5 to 10 minutes. Each spectrum is an average of 5-10 scans with scan time of 60 seconds.

### The universal Point Accumulation Imaging in Nanoscale Topography (uPAINT)

uPAINT experiments were performed as described earlier (*31*). Hippocampal neurons from C57BL6/J and *DDHD2^−/−^* were transfected with VAMP2–pHluorin (a kind gift from Dr. Volker Haucke, Leibniz-Forschungsinstitut für Molekulare Pharmakologie Berlin) at DIV14. To track VAMP2–pHluorin mobility at the presynapses in resting and stimulated conditions, we diluted anti-GFP Atto647N nanobodies (Synaptic Systems, cat. no. N0301-At647N-S) to final concentration of 100 pM in low K^+^ or high K^+^ buffer. On DIV20-22, cultured hippocampal neurons expressing VAMP2–pHluorin were washed once with low K^+^ buffer and then imaged in low K^+^ buffer containing 100 pM anti-GFP Atto647N nanobodies for 10,000 frames (50 Hz, 20 ms exposure). The unbound anti-GFP Atto647N nanobodies were then washed off with 8-10 mL of low K^+^ buffer, and then subsequently imaged in high K^+^ containing 100 pM anti-GFP Atto647N nanobodies for 10,000 frames (50 Hz, 20 ms exposure). Imaging was done at 37 °C on a Roper Scientific Ring-TIRF microscope equipped with an iLas2 double laser illuminator (Roper Scientific, FL, USA), a Nikon CFI Apo TIRF 100/1.49 N.A. oil-immersion objective (Nikon Instrument, NY, USA) and a Perfect Focus System (Nikon Instruments, NY, USA). Imaging was performed using two Evolve512 delta EMCCD cameras (Photometrics, AZ, USA) mounted on a TwinCam LS Image Splitter (Cairn Research, UK), a quadruple beam splitter (ZT405/488/561/647rpc, Chroma Technology, VT, USA) and a QUAD emission filter (ZET405/488/561/640 m, Chroma Technology, VT, USA).

### Single-particle tracking photoactivated localization microscopy (sptPALM)

Hippocampal neurons from C57BL6/J and *DDHD2^−/−^* were transfected with pmEOS2-C1-ERGIC53 (ERGIC-53-mEos2) at DIV14. On DIV20-22, time-lapse movies (10,000 frames) were acquired at 50 Hz using Metamorph software as described above. The 405-nm laser was used to photo-convert ERGIc-53-mEos2, and the 561-nm laser was used simultaneously for excitation and bleaching of the resulting photo-converted single molecules. To isolate the mEos2 signal from autofluorescence and background signals, a double-beam splitter (LF488/561-A-000, Semrock, NY, USA) and a double-band emitter (FF01-523/610-25, Semrock, NY, USA) were used. To spatially distinguish and temporally separate the stochastically activated molecules during acquisition, the 405-nm laser was used between 1.5% and 5% of the initial laser power (100 mW Vortran Laser Technology), and the 561-nm laser was used at 80% of the initial laser power (150 mW Cobolt Jive).

### Single particle tracking

PALMTracer (*52–54*) in Metamorph software (MetaMorph Microscopy Automation and Image Analysis Software, v7.7.8; Molecular Devices) was used to obtain the mean square displacement (MSD) and diffusion coefficient (D; μm^2^ s^−1^) values. Tracks shorter than eight frames were excluded from the analysis to minimize nonspecific background. The Log_10_D immobile and mobile fraction distributions were calculated as previously described (*55*), setting the displacement threshold to 0.03 μm^2^ s^−1^ (*i.e*. Log_10_D = −1.45 when [D] = μm^2^ s^−1^). The mobile to immobile (M/IMM) ratio was determined based on the frequency distribution of the diffusion coefficients (Log_10_D) of immobile (Log_10_D ≤ −1.45) and mobile (Log_10_D > −1.45) molecules (the immobile fraction of molecules represents LifeAct-mEos2 molecules for which the displacement within 4 frames was below the spatial detection limit of our methods, 106 nm). The area under the MSD curve (AUC) was calculated in Prism 9 for macOS version 9.1.1 (GraphPad Prism 9 for macOS). The super-resolved image color-coding was done as previously described(*43*) using ImageJ/Fiji (2.0.0-rc-43/1.50e; National Institutes of Health), with each coloured pixel in the average intensity maps indicating the localization of an individual molecule (bar: 8 to 0, high to low density), the color-coded pixels in the average diffusion coefficient map presenting an average value for each single-molecule track at the site of localization (bar: Log_10_ 1 to −5, high to low mobility), and the color-coding of the track maps representing the detection time point (bar: 0–10,000 frame acquisition) during acquisition.

### Quantification and statistical analysis

Statistical tests were conducted in Prism 9 for macOS version 9.1.1. The normality of the data was tested with Kolmogorov–Smirnov tests, and nonparametric tests and parametric tests were used to compare two independent groups (Mann–Whitney test or t test) and multiple groups (Kruskal–Wallis test or ordinary 2way ANOVA with Tukey post hoc multiple comparison test). Dot plots indicate average ± SEM from individual acquisitions or ROIs unless otherwise stated.

---

**Fig. S1.**
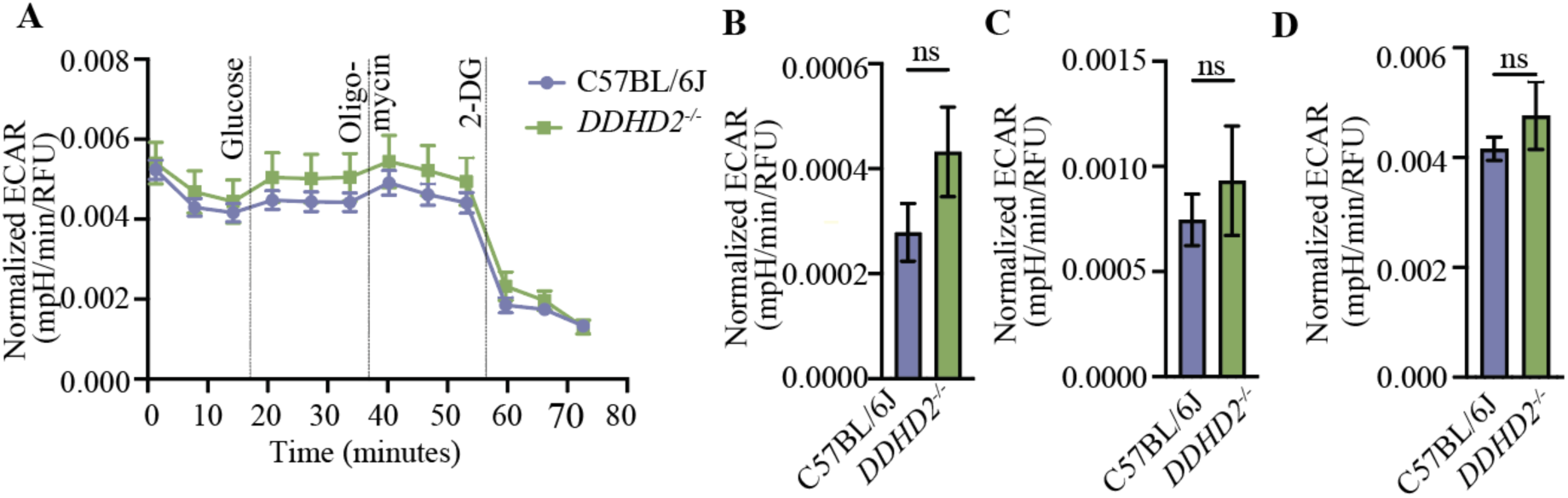
Extracellular acidification rate remains largely unchanged following the loss of DDHD2. (**A**) Seahorse XF measurement of extracellular acidification rate (ECAR) in cultured E16 neurons from C57BL/6J and *DDHD2^−/−^.* Injection of glucose, oligomycin (an ATP synthase inhibitor), and 2-deoxy-D-glucose (2-DG; a glucose analog which inhibits glycolysis through competitive binding of glucose hexokinase, the first enzyme in the glycolytic pathway) are indicated. Quantification of ECAR (**B**) basal glycolysis, (**C**) glycolytic capacity, (**D**) non-glycolytic acidification in indicated conditions. Data are the mean ± sem, student’s t test, non-significant (ns), N=2 independent experiments in each condition.

**Fig. S2.**
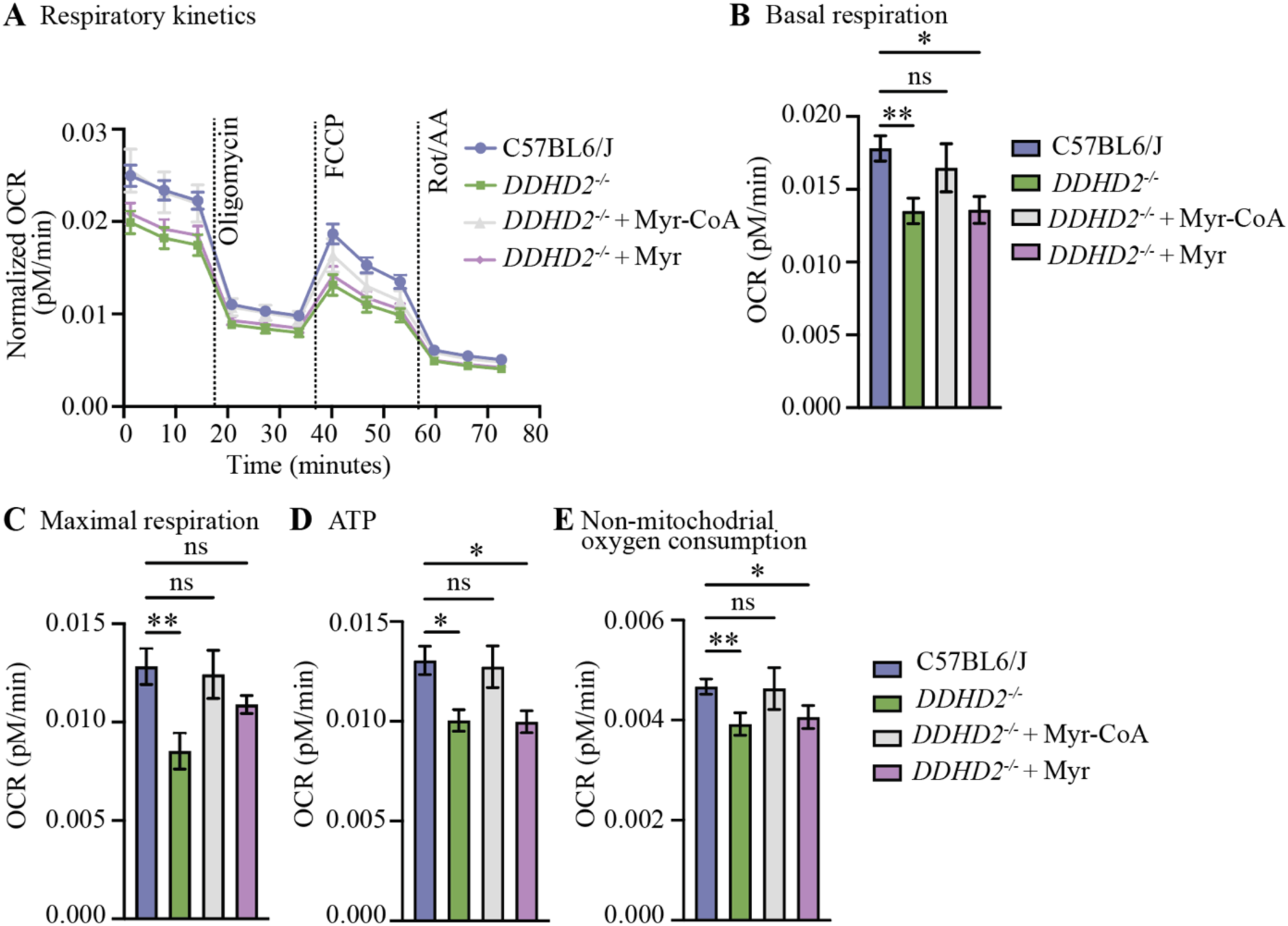
Without CoA-conjugation, myristic acid does not rescue the mitochondrial respiratory function in *DDHD2^−/−^* neurons. (**A**) Seahorse XF measurement of oxygen consumption rate (OCR), (**B**) basal respiration, (**C**) maximal respiration, (**D**) ATP production, and (**E**) non-mitochondrial oxygen consumption in cultured neurons from C57BL6/J, *DDHD2^−/−^* and *DDHD2^−/−^* treated with 1 µM Myr-CoA or 1 µM myristic acid for 48 h. Data are the mean ± sem, N=1 with 14-76 technical replicates, Kruskal-Wallis multiple comparisons test, non-significant (ns), *P<0.05, **p<0.01.

**Fig. S3.**
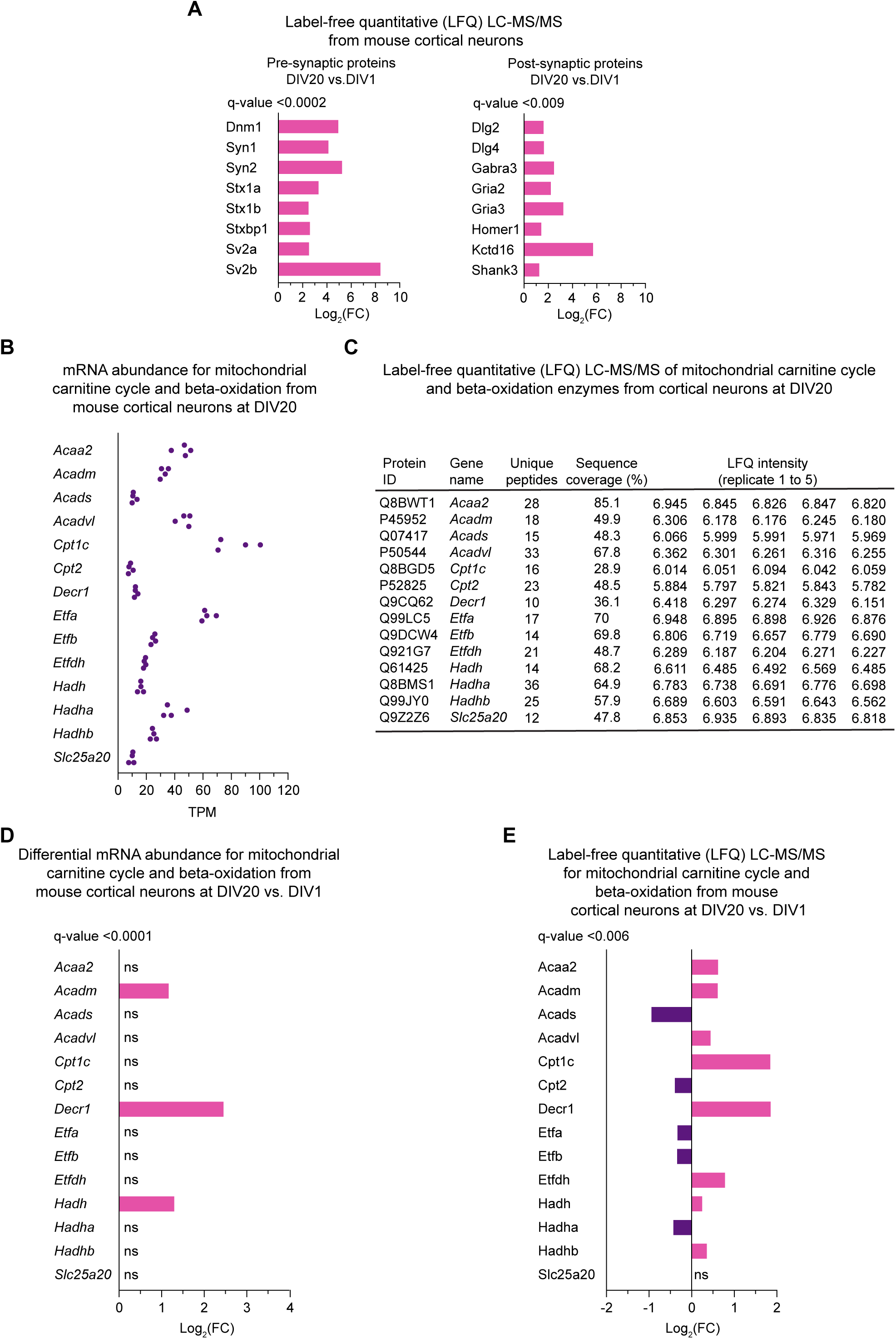
Mitochondrial carnitine cycle and β-oxidation enzymes expression in cultured cortical neurons. (**A**) Label-free quantitative (LFQ) LC-MS/MS expression analysis of pre-synaptic and post-synaptic proteins from cultured mouse cortical neurons at DIV20 vs. DIV1. (**B**) Total RNAseq expression of mRNAs encoding enzymes of the mitochondrial carnitine cycle and β-oxidation at DIV20 from cultured mouse cortical neurons. TPM, transcript per million. (**C**) Label-free quantitative (LFQ) LC-MS/MS expression analysis of the enzymes of the mitochondrial carnitine cycle and beta-oxidation at DIV20 from cultured mouse cortical neurons. (**D**) DESeq2 comparison of gene expression changes for mRNAs encoding enzymes of the mitochondrial carnitine cycle and beta-oxidation at DIV20 vs. DIV1 from cultured mouse cortical neurons. (**E**) Label-free quantitative (LFQ) LC-MS/MS expression analysis for proteins of the mitochondrial carnitine cycle and beta-oxidation at DIV20 vs. DIV1 from cultured mouse cortical neurons. N=5 independent replicates for each condition.

**Fig. S4.**
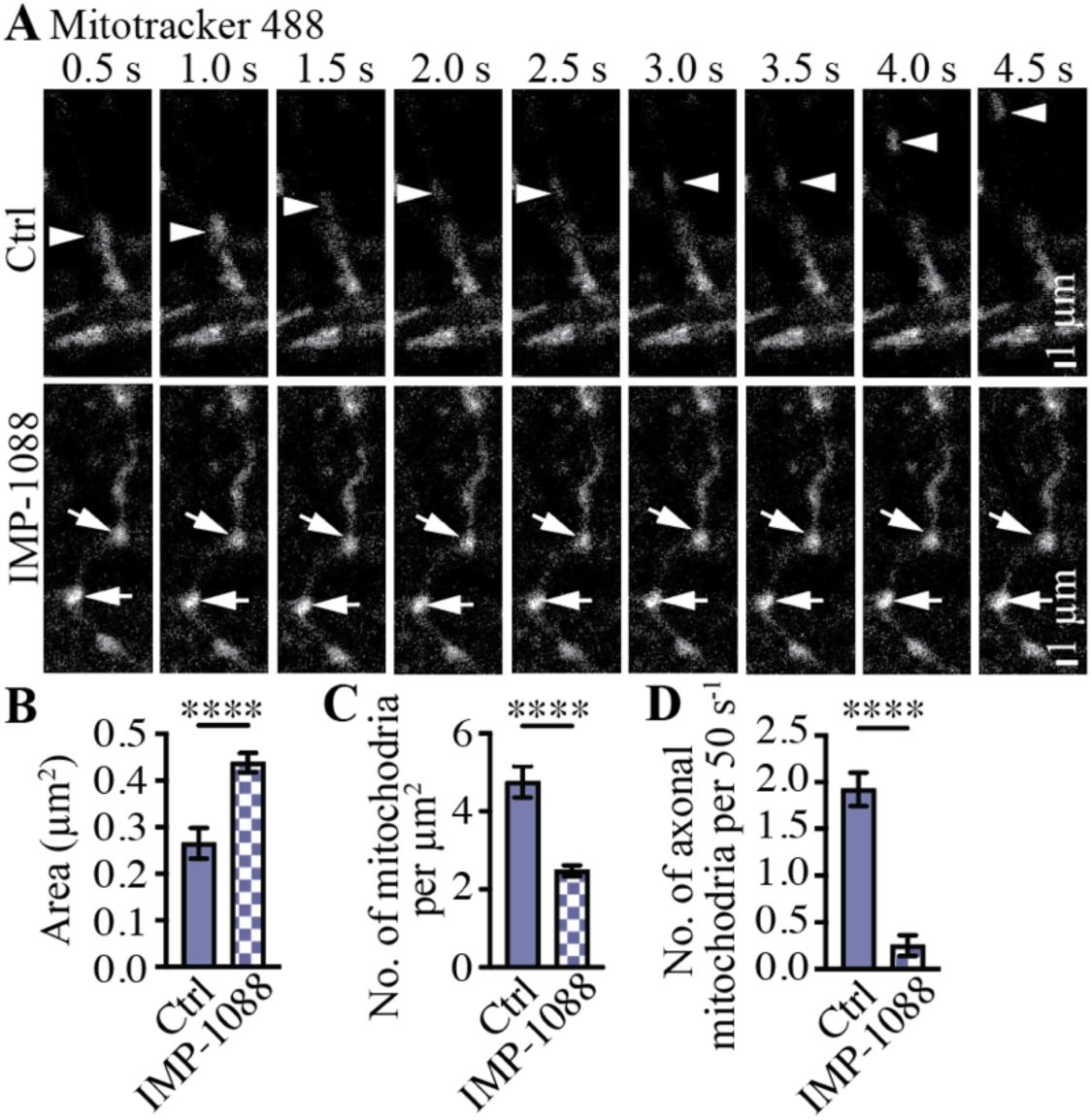
Live cell imaging reveals that an inhibition of N-myristoylation affects the structure and dynamics of mitochondria in neurons. (**A**) Representative confocal images of cultured control (vehicle DMSO) and IMP-1088 (1 μM for 48h) treated C57BL6/J neurons and imaged using MitoTracker dye. Arrowheads point to a mobile and arrows to stalled mitochondria in the synapses. Quantification of (**B**) mitochondrial area (μm^2^), (**C**) number of mitochondria per μm^2^ and (**D**) number of axonal transported mitochondria. Data are the mean ± sem, student’s t test, ****p<0.0001, N=2 independent experiments in both conditions.

**Fig. S5.**
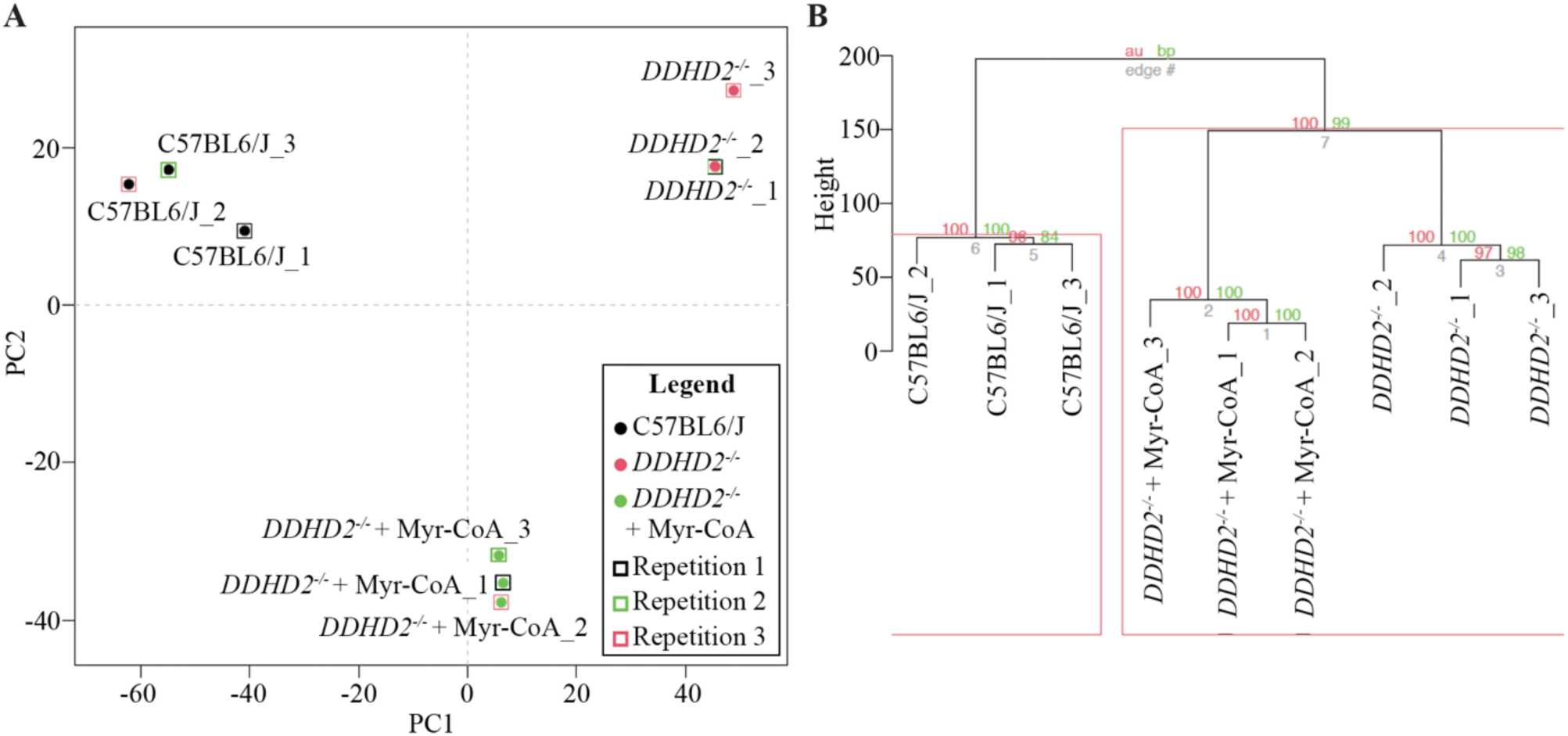
Mass spectrometry analysis of C57BL6/J, *DDHD2^−/−^* and *DDHD2^−/−^* treated with 1 µM Myr-CoA for 48 h. (**A**) Principal component analysis. The first two principal components that explained the most amount of variation is shown (x and y-axis). Samples can be observed to cluster according to their experimental group. (**B**) Hierarchical clustering. Samples cluster according to their experimental group (within each of the tree-like - dendrograms). Values on branches and the red box, represent probabilities of observing the clustering after 100 bootstraps. N = 3 independent experiments in each group.

**Fig. S6.**
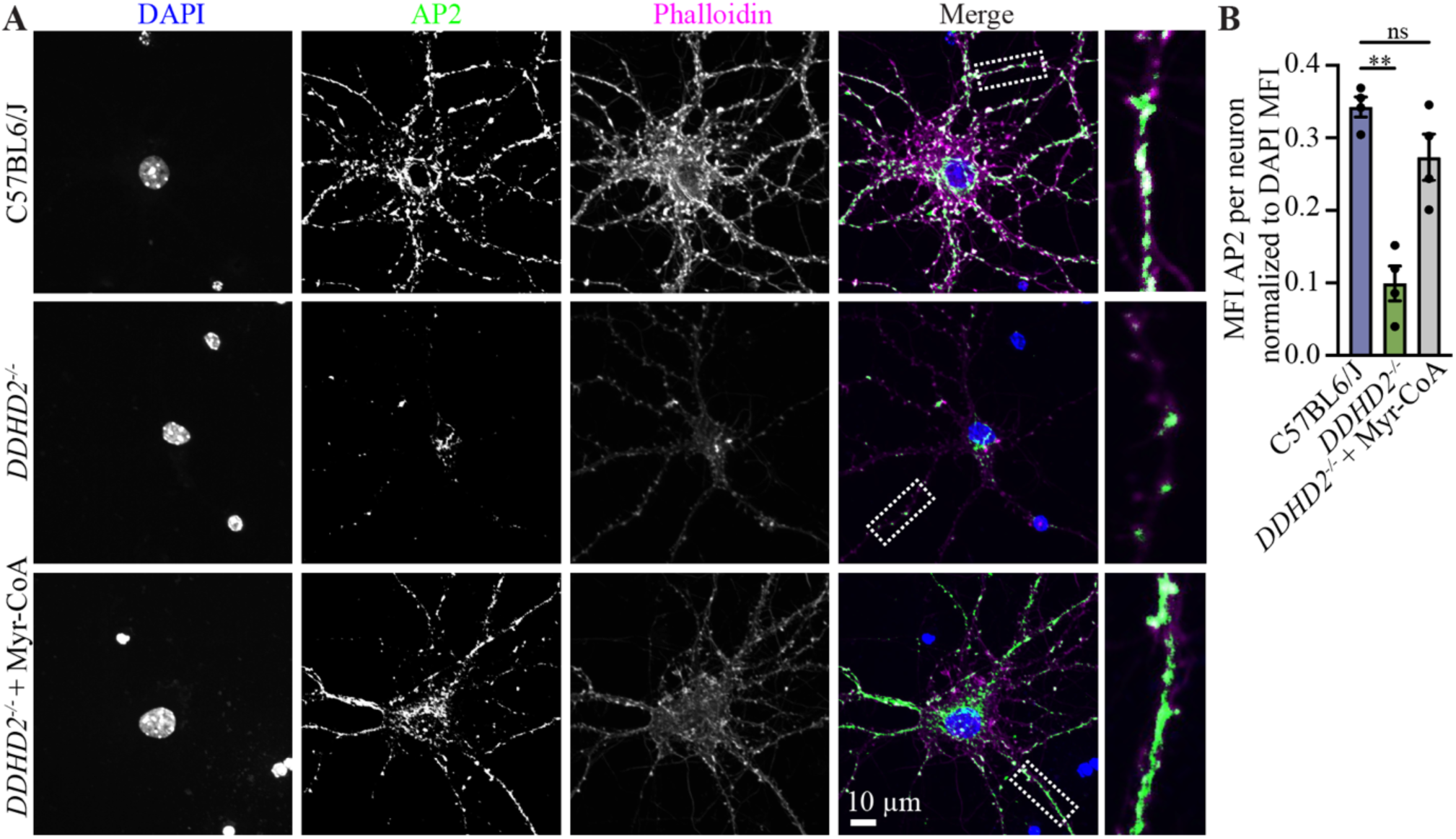
Mislocalization of clathrin adaptor AP-2 associated with the loss of DDHD2 is significantly rescued with Myr-CoA supplementation. (**A**) Representative maximum projection confocal images of cultured hippocampal neurons C57BL6/J, *DDHD2^−/−^*, and *DDHD2^−/−^*+1 µM Myr-CoA for 48 h, immunostained for endogenous AP-2 (green). DAPI (nucleus, blue) and Phalloidin (actin, magenta) staining are shown for reference. Boxed areas are shown magnified on right. (**B**) Quantification of mean fluorescent intensity (MFI) of AP-2 per neuron, normalized to DAPI MFI in indicated conditions. Data are the mean ± sem (n=3). Kruskal-Wallis multiple comparisons test. Ns, non-significant, **p<0.01.

**Fig. S7.**
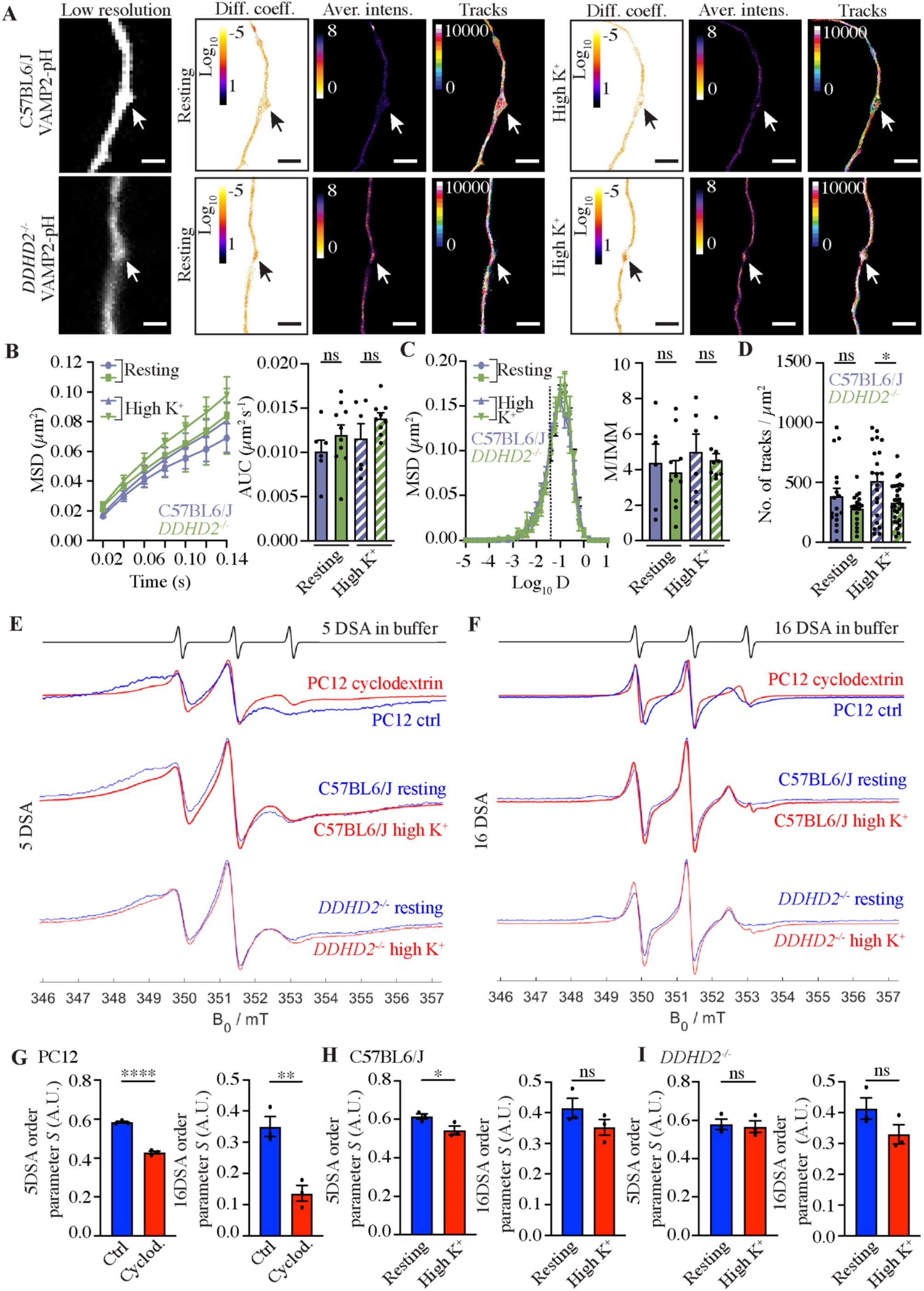
Single molecule imaging and neuronal plasma membrane fluidity measurement in C57BL6/J and *DDHD2^−/−^* neurons. (**B**) uPAINT super-resolution imaging VAMP2-pHluorin (VAMP2-pH) bound Atto647N-nanobodies in cultured C57BL6/J and *DDHD2^−/−^* neurons in resting condition (low K^+^) and following high K^+^ stimulation. Low resolution images of green fluorescence of the VAMP2-pHluorin, along with super-resolved diffusion coefficient (bar: Log_10_ 1 to −5, high to low mobility), average intensity (bar: 8 to 0, high to low density), and single-molecule trajectory (bar: 0–10,000 frame acquisition) maps are show. Arrows point to presynapses. Quantification of single-molecule mobility of VAMP2-pH/Atto647N-nanobodies in C57BL6/J and *DDHD2^−/−^* neurons shown as (**B**) mean square displacement (MSD) and area under the MSD curve, (**C**) frequency distribution of log_10_ diffusion coefficients ([D] = μm^2^ s^−1^), and mobile to immobile ratio of diffusion coefficient frequency distributions (immobile Log_10_D ≤ −1.45 and mobile Log_10_D > −1.45) in indicated conditions. (**D**) Quantification of single-molecule numbers of VAMP2-pH/Atto647N-nanobodies per µm^2^ in presynapses in C57BL6/J and *DDHD2^−/−^* neurons in resting and high K^+^ stimulated conditions. (**E-I**) X-band (9.852) CW EPR_spectra_ recorded from E16 C57BL6/J and *DDHD2−/−* neurons at DIV21-22 in resting condition (non-stimulated) and following a 5-min high K^+^ stimulation using a microwave power of 16 mW and a modulation amplitude of (**E**) 0.3 mT (DSA5) and (**F**) 0.1 mT (DSA16). PC12 neurosecretory cells treated with 1 µM cyclodextrin for 30 min or DMSO (vehicle media; Ctrl) are shown as a positive control. Quantification of 0.3 mT (DSA5) and 0.1 mT (DSA16) membrane order parameter *S* (A.U. arbitrary units) in (**G**) PC12 cells, (**H**) C57BL6/J and (**I**) *DDHD2^−/−^* neurons in indicated conditions. Data are the mean ± sem, Kruskal-Wallis multiple comparison test, N = 3 independent experiments, non-significant (ns), *p<0.05, **p<0.01, ***p<0.001, and ****p<0.0001.

**Table 1.** Comparative proteomic analysis of cortical neurons from wild type NMRI mice at DIV1 and DIV20.

**Table 2.** mRNA expression analysis of cortical neurons from wild type NMRI mice.

**Table 3.** Proteomic analysis of hippocampal neurons from wild type C57BL6/J, *DDHD2^−/−^* and *DDHD2^−/−^* supplemented with 1 µM Myr-CoA for 48 h at DIV21-22.

**Movie 1. Live imaging of C57BL6/J hippocampal neuron stained with Mitotracer-488.** The neurons were treated with DMSO for 48 h (vehicle control). The confocal imaging was done at 2 frames s^−1^, and the playback is 50 frames s^−1.^

**Movie 2. Live imaging of C57BL6/J hippocampal neuron treated with IMP-1088 and stained with Mitotracer-488.** The neurons were treated with 1 µM IMP-1088 for 48 h. The confocal imaging was done at 2 frames s^−1^, and the playback is 50 frames s^−1.^

